# Axonal ER Ca^2+^ Release Enhances Miniature, but Reduces Activity-Dependent Glutamate Release in a Huntington Disease Model

**DOI:** 10.1101/2020.01.31.929299

**Authors:** James P. Mackay, Amy I. Smith-Dijak, Ellen T. Koch, Peng Zhang, Evan Fung, Wissam B. Nassrallah, Caodu Buren, Mandi Schmidt, Michael R. Hayden, Lynn A. Raymond

**Affiliations:** Department of Psychiatry, Djavad Mowafaghian Centre for Brain Health, University of British Columbia, Vancouver, British Columbia, Canada; Graduate Program in Neuroscience, University of British Columbia, Vancouver, British Columbia, Canada; MD/PhD Program, University of British Columbia, Vancouver, British Columbia, Canada; Centre for Molecular Medicine and Therapeutics, BC Children’s Hospital Research Institute, University of British Columbia, Vancouver, British Columbia, Canada

## Abstract

Action potential-independent (miniature) neurotransmission occurs at all chemical synapses, but remains poorly understood, particularly in pathological contexts. Spontaneous release of Ca^2+^ from the axonal endoplasmic reticulum (ER) is thought to facilitated miniature neurotransmission, and aberrant ER Ca^2+^ handling is notably implicated in the progression of Huntington’s disease (HD) and other neurodegenerative diseases. Here, we report elevated glutamate-mediated miniature synaptic event frequencies in YAC128 (HD-model) cortical neurons, which pharmacological experiments suggest is mediated by enhanced spontaneous ER Ca^2+^ release. Calcium imaging using an axon-localized sensor revealed slow action potential (AP)-independent axonal Ca^2+^ waves, which were more common in YAC128 cortical neurons. Conversely, spontaneous axonal ER Ca^2+^ release was associated with reduced AP-dependent axonal Ca^2+^ events and consequent glutamate release. Together, our results suggest spontaneous release of axonal ER Ca^2+^ stores oppositely regulates activity-dependent and -independent neurotransmitter release in HD, with potential implications for the fidelity and plasticity of cortical excitatory signaling.

## Introduction

Huntington’s disease (HD) is a fatal, autosomal dominantly-inherited neurodegenerative disorder caused by a polyglutamine-encoding CAG repeat-expansion (>35 repeats) in exon-1 of the huntingtin gene (“A novel gene containing a trinucleotide repeat that is expanded and unstable on Huntington’s disease chromosomes. The Huntington’s Disease Collaborative Research Group.,” 1993). Disease onset is typically in middle age, characterized by progressively disordered movement and declining cognition (Bachoud-Lévi et al., 2019). Although the mutant huntingtin protein (mHTT) is widely expressed, GABAergic spiny projection neurons (SPN)s of the striatum and pyramidal neurons of the cerebral cortex show the most severe degeneration in HD (Graveland, Williams, & DiFiglia, 1985; Vonsattel et al., 1985). Cortical glutamatergic afferents extensively innervate striatal SPNs and dysfunction at these synapses is thought to precede overt neuron loss in HD (Raymond et al., 2011).

Glutamate-mediated toxicity was initially proposed to contribute to pathogenesis in HD based on studies showing that striatal injections of glutamate receptor agonists largely recapitulate HD pathology in animals (Beal et al., 1986; Hantraye, Riche, Maziere, & Isacson, 1990). More recently, studies in SPNs from transgenic and knock-in mouse models of HD demonstrate enhanced extrasynaptic N-Methyl-D-Aspartate receptor (NMDAR) surface expression and function that may explain, in part, increased susceptibility to excitotoxic challenges (Botelho et al., 2014; Fan, Fernandes, Zhang, Hayden, & Raymond, 2007; Kovalenko et al., 2018; Milnerwood et al., 2010; Plotkin et al., 2014; Zeron et al., 2002). Cell stress/death signaling mediated by elevated levels of extrasynaptic NMDARs in HD SPNs may be further exacerbated by reduced glutamate transporter function (Miller et al., 2008), although recent studies show glutamate clearance after synaptic release is normal or accelerated in striatal and cortical brain slice (Parsons et al., 2016). Additionally, some studies show synaptic glutamate release from cortical afferents is altered in HD mouse models, possibly contributing to excitotoxicity and synaptic dysfunction (Cepeda et al., 2003; Joshi et al., 2009; Raymond et al., 2011). However, the direction of this effect appears to be model and disease-stage dependent; relative to wildtype (WT), glutamate release is typically enhanced early, but reduced with disease progression in HD mice (Joshi et al., 2009).

Mutant HTT (mHTT) directly interacts with endoplasmic reticulum (ER) type-1 inositol (1,4,5)-triphosphate receptors (IP3Rs) sensitizing their Ca^2+^ release in response to IP3 (Tang et al., 2003); blocking this interaction normalizes IP3-induced ER Ca^2+^ release *in vitro* and improves behavioral outcomes in HD-model mice (Tang, Guo, Wang, Chen, & Bezprozvanny, 2009). Furthermore, evidence suggests Ryanodine receptors, which are ER-localized Ca^2+^ channels that mediate Ca^2+^-induced Ca^2+^ release, are constitutively leaky in HD mouse models (Suzuki, Nagai, Wada, & Koike, 2012). Although these studies focused on the soma and dendrites of SPNs at the postsynaptic side of cortical-striatal synapses, ER is found in all neuronal compartments, including presynaptic terminals, where its Ca^2+^ stores, when released, modulate neurotransmission (Emptage, Reid, & Fine, 2001; Llano et al., 2000). However, it is unknown whether presynaptic ER Ca^2+^ handling is also dysfunctional in HD and if such a process contributes to altered glutamate release from cortical synaptic terminals.

Here, we have investigated whether glutamate release is altered, and if ER Ca^2+^ dysregulation contributes to aberrant function, in presynaptic terminals of cortical pyramidal neurons from premanifest HD-model mice expressing full-length mHTT with 128 CAG-repeats in a yeast artificial chromosome (YAC128). Our results from neuronal cultures and acute cortical-striatal brain slices demonstrate a shift in balance favoring increased action potential (AP)-independent (miniature) vs. AP-dependent glutamate release, and that altered ER Ca^2+^ release contributes to this change. This may have important previously unidentified disease implications, given the distinct physiologically relevant signaling potentially mediated by miniature neurotransmission (Fishbein & Segal, 2007; Kavalali, 2015). Ultimately our data contribute to growing evidence for cortex and striatal synaptic dysfunction in HD, and support a key role for ER Ca^2+^ dysregulation in this pathophysiology.

## Results

### Mini EPSC frequencies are elevated in YAC128 cortical cultures at early time points

Our group previously recorded miniature excitatory post synaptic currents (mEPSCs) from striatal spiny projection neurons (SPNs) co-cultured with cortical neurons from prenatal WT or yeast artificial chromosome (YAC128) mice. YAC128 SPNs showed elevated mEPSC frequencies, compared to WT, at day in vitro (DIV) 14 – a time point when SPN dendritic arborization patterns and spine numbers were similar between genotypes (Buren, Parsons, Smith-Dijak, & Raymond, 2016). At DIV21, YAC128 SPNs showed a significant reduction in total dendritic length, and therefore reduced total excitatory synapse numbers, compared to WT. Despite this, mEPSC frequencies onto WT and YAC128 SPNs were comparable at DIV21.

The above results suggested a higher rate of action potential-independent glutamate release from YAC128 presynaptic cortical pyramidal neuron (CPN) terminals onto individual SPN synapses. Here, we capitalize on a relatively simpler culture preparation containing only cortex-derived neurons and YAC128 mouse-derived brain slices to mechanistically dissect how mutant huntingtin protein expression affects action potential-independent and -dependent glutamate release from CPN terminals.

To establish whether miniature glutamate release is enhanced at YAC128 CPN terminals targeting other CPNs, we first established WT and YAC128 CPN mEPSC parameters in cortical cultures at various DIV ages (7, 14, 18 and 21). In general, both genotypes showed increased mEPSC frequencies with culture time. However, YAC128 CPN mEPSC frequencies were consistently higher than age-matched WT controls after DIV7 and up until DIV21, at which point mEPSC frequencies became comparable between genotypes **(Figure 1)**, as was also the case in the co-culture model (Buren et al., 2016). The greatest genotype mEPSC frequency difference occurred at DIV14, when mean YAC128 CPN frequencies were more than double that of WT: 10.83 ± 1.80 Hz vs 4.52 ± 0.60 Hz, respectively **(Figure 1D, E)**. YAC128 mEPSC frequencies remained significantly higher than WT at DIV18: 14.07 ± 1.54 Hz vs 9.92 ± 1.54 Hz, respectively **(Figure 1G, H)**. Age matched YAC128 and WT CPNs showed similar mEPSC amplitudes across all DIV time points examined **(Figure 1 C, F, I and L)**. We chose to perform subsequent experiments probing the mechanistic details underlying increased cortical mini glutamate release in DIV18 cortical monocultures, since this was the most mature culture stage at which mEPSC frequencies were elevated in YAC128 CPNs.

**Fig 1:**
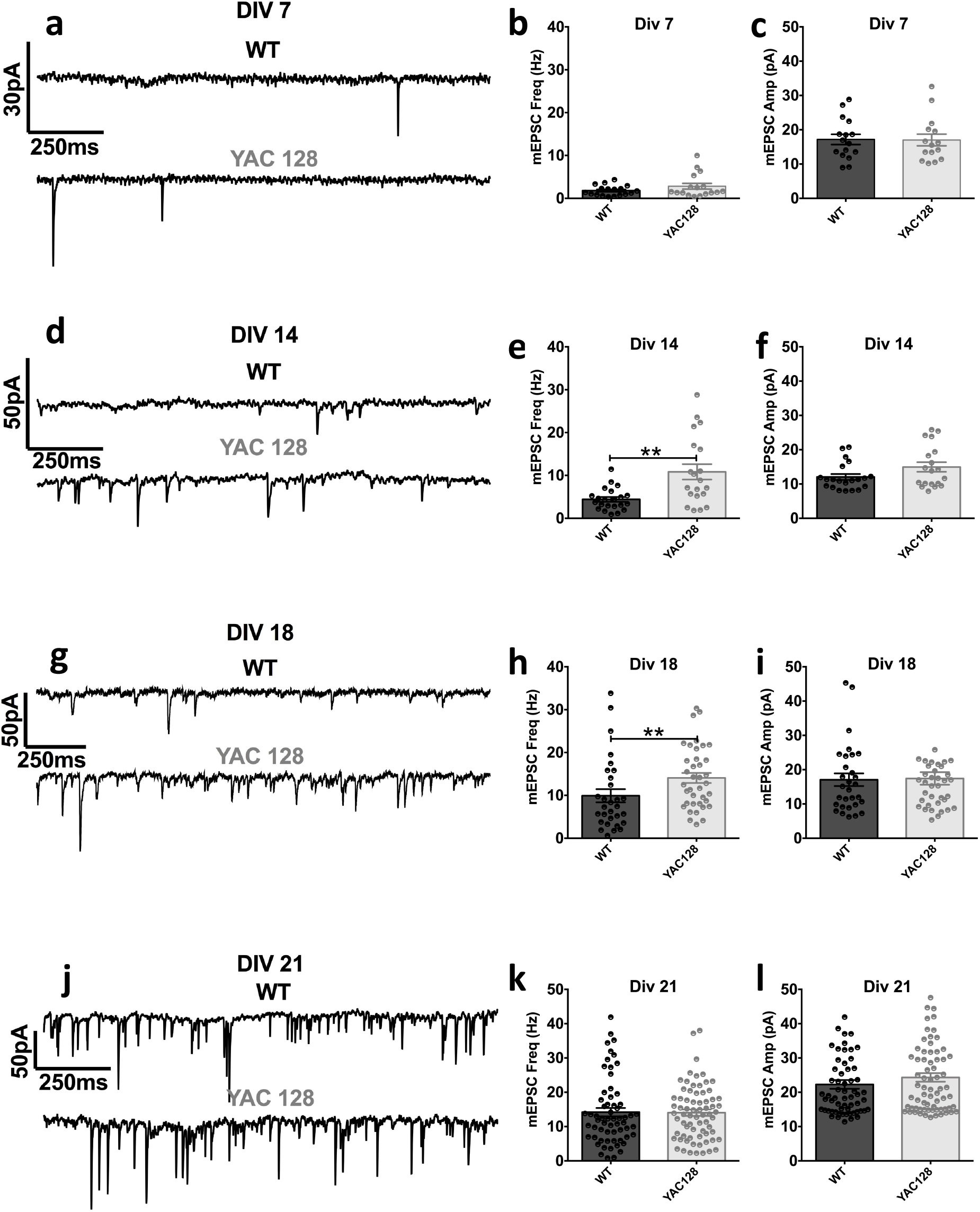
YAC128 cortical cultures show elevated miniature EPSC frequencies at early DIV time points. Representative traces (**A, D, G, J**) and population data for mEPSCs (frequency in **B, E, H, K**; amplitude in **C, F, I, L**) recorded from WT and YAC128 CPNs. All recordings were made in voltage clamp with a holding potential of -70mV in solution containing TTX (500 nM) and PTX (50 μM). **A – C:** Recordings made from CPNs in DIV-7 cultures. Mean mEPSC frequency (**B**) was 1.8 ± 0.3 Hz (n=18; 3 cultures) in WT CPNs and 2.8 ± 0.7 Hz (n=17; 4 cultures) in YAC CPNs; this difference was not statistically significantly [t(33)=1.449; p=0.1567; Student’s unpaired-t test]. Mean mEPSC amplitudes (**C**) were similar in WT and YAC128 cultures: 17.2 ± 1.5 pA (n=18; 3 cultures) and 17.0 ± 1.7 pA (n=17; 4 cultures) respectively [t(33)=0.06936; p=0.9452; Student’s unpaired-t test]. **D - F:** Recordings were made from CPNs in DIV-14 cortical cultures. Mean mEPSC frequency (**E)** was significantly higher in YAC128 CPNs [10.8 ± 1.8 Hz (n=20; 6 cultures)] compared to WT CPNs [4.4 ± 2.6 Hz (n=22; 4 cultures)], [t(40)=3.555; p=0.0010; Student’s unpaired-t test]. Mean CPN mEPSC amplitudes (**F**) were not significantly different in WT and YAC128 cultures: 12.1 ± 0.8 pA (n=22; 4 cultures) and 15.0 ± 1.4 pA (n=20; 6 cultures), respectively [t(40)=1.807; p=0.0786; Student’s unpaired-t test]. **G – I:** Recordings were made from CPNs in DIV-18 cortical cultures. Mean CPN mEPSC frequency (**H**) was significantly higher in YAC128 cortical cultures [14.1 ± 1.2 Hz (n=38; 10 cultures)] compared to WT cultures [9.9 ± 1.5 Hz (n=30; 15 cultures)], [p<0.0059 (exact); Mann Whitney test]. Mean CPN mEPSC amplitudes (**I**) were similar in WT and YAC128 cultures: 17.0 ± 1.8 pA (n=30; 15 cultures) and 17.4 ± 1.8 pA (n=38; 10 cultures) respectively [p=0.8643 (exact); Mann Whitney test]. **J – L:** Recordings were made from CPNs in DIV-21 cortical cultures. Mean CPN mEPSC frequency (**K**) was similar in WT and YAC128 cultures: 14.2 ± 1.2 Hz (n=62; 17 cultures) and 14.0 ± 0.9 Hz (n=71, 23 cultures), respectively [p=0.5626 (exact); Mann Whitney test]. Mean CPN mEPSC amplitudes (**L**) were similar in WT and YAC128 cultures: 22.3 ± 1.3 pA (n=62; 17 cultures) and 24.3 ± 1.2 pA (n=71; 23 cultures), respectively [p=0.2143 (exact); Mann Whitney test].

### Synapse numbers and dendritic complexity are similar in WT and YAC128 cortical pyramidal neurons at DIV 18

The higher mEPSC frequencies seen in DIV18 YAC128 CPNs suggest an increased presynaptic glutamate release probability. However, relative differences in synapse numbers could also account for this finding. To estimate numbers of synapses, we first expressed cytosolic GFP in a small proportion of neurons in WT and YAC128 cortical cultures and imaged full CPN dendritic arbors at DIV18. Sholl analysis revealed similar arborization patterns and total dendritic length in WT and YAC128 CPNs **(Supplemental Figure 1A, C)**. In separate cultures, we expressed an internal GFP-tagged anti-PSD95 antibody (Gross et al., 2013) in a subset of neurons and immuno-stained for VGlut1, to identify glutamatergic terminals, and the GluA2 AMPA receptor subunit, to identify functional synapses. Functional synapse numbers, defined as GFP-labeled PSD95 puncta colocalized with VGlut1 and GluA2 immunofluorescent-labeled puncta, were not significantly different between DIV18 WT and YAC128 CPNs, although there was a trend towards lower synapse density in YAC128 CPNs **(Supplemental Figure 1E, F)**. Together with the above data showing increased mEPSC frequency in YAC128 CPNs at DIV18 **(Figure 1)**, these results point to an increase in miniature vesicular glutamate release from cortical terminals in YAC128 cultures.

### Releasing ER calcium with low dose ryanodine or caffeine increases the miniature EPSC frequency in WT, but not YAC128 cultures

Studies using mouse models suggest Ca^2+^ release from ER stores is aberrant in HD due to increased IP3 and ryanodine receptor activity (Suzuki et al., 2012; Tang et al., 2003). Although effects of mHTT on the presynaptic ER have not been specifically studied, we hypothesized a presynaptic ER leak elevates cytosolic Ca^2+^, thereby mediating the increased miniature glutamate release seen in our YAC128 cultures **(Figure 2A, B)**. To test this hypothesis, we first recorded mEPSCs before and during local application of low dose (5 μM) ryanodine to cultured CPNs; 5 μM ryanodine releases ER Ca^2+^ by opening ryanodine receptors (Meissner, 2017). In WT cultures, 5 μM ryanodine nearly doubled the CPN mEPSC frequency: from 6.74 ± 1.17 Hz to 11.91 ± 2.36 Hz [Students’ paired T-test; p=0.0280; n=13 cells] **(Figure 2C - E)**. In YAC128 cultures, however, ryanodine (5 μM) did not significantly alter the CPN mEPSC frequency: 11.98 ± 1.66 Hz vs 13.89 ± 2.01 Hz - under control conditions and in ryanodine (5 μM) respectively **(Figure 2F - H)**. A Comparison of the percent change in mEPSC frequency following ryanodine (5 μM) revealed a significantly greater response in WT than in YAC128 CPNs [31.7 ± 7.5 % (n=13) vs 5.2 ± 9.5 % (n=15) in WT and YAC128 CPNs respectively (Students’ unpaired T-test; p=0.0418)]. We next used caffeine (1 mM) as an alternative means of agonizing ryanodine receptors; this more prominently increased the mEPSC frequency in WT cultures, but otherwise produced similar results **(Supplemental Figure 2)**.

**Fig 2:**
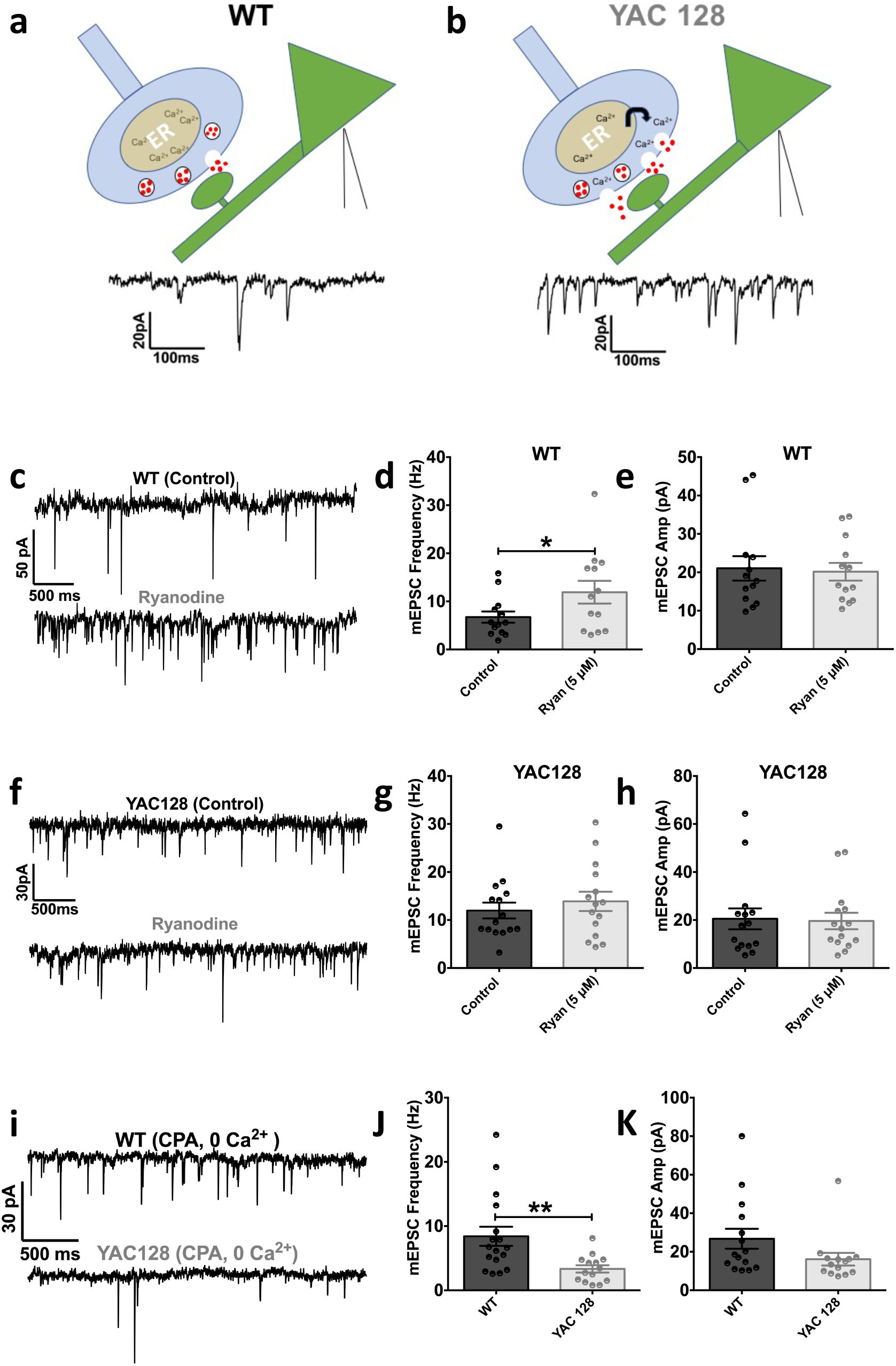
Spontaneous release of ER Ca^2+^ elevates YAC128 pyramidal neuron miniature EPSC frequencies. **A and B - Hypothetical model (tested below) A**. We hypothesize, in WT cultures, spontaneous Ca^2+^ release from ER stores in presynaptic cortical terminals only minimally augments action potential-independent (mini) glutamate release. **B**. Conversely, in YAC128 cultures, we hypothesize spontaneous Ca^2+^ release from presynaptic ER stores is increased, and that this elevates Ca^2+^-dependent, mini glutamate release. **C – H:** Voltage clamp recordings were made at -70 mV from DIV-18 cultured WT (**C – E**) and YAC128 CPNs (**F – H**) in the presence of TTX (500 nM) and PTX (50 μM), under control conditions and during subsequent local ryanodine (5 μM) application. Note that in the representative traces from the WT CPN (**C**), ryanodine (5 μM) substantially increased the mEPSC frequency, whereas the drug had little effect on the mEPSC frequency in the representative YAC128 CPN (**F**). Quantifying the population data revealed that ryanodine (5 μM) significantly increased the mean mEPSC frequency in WT cultured CPNs (**D**) from 6.7 ± 1.2 Hz to 11.9 ± 2.4 Hz, [t(12)=2.50; p=0.0280; n=13; 8 cultures; Student’s paired-t test]. Ryanodine (5 μM) did not significantly affect the mean WT CPN mEPSC amplitude (**E**) (control: 21.0 ± 3.2 pA, ryanodine: 20.2 ± 2.3 pA), [t(12)=0.65; p=0.5278; n=13; 8 cultures; Student’s paired-t test]. In YAC128 cultures, ryanodine (5 μM) did not significantly affect the mean CPN mEPSC frequency (**G**) (control: 12.0 ± 1.7 Hz, ryanodine: 13.9 ± 2.0 Hz), [t(14)=1.65; p=0.1206; n=15; 4 cultures; Student’s paired-t test]. Ryanodine (5 μM) also did not significantly affect the mean CPN mEPSC amplitude (**H**) (control: 20.5 ± 4.4 pA, ryanodine: 19.6 ± 3.4 pA), [t(14)=0.71; p=0.4904; n=15; 4 cultures; Student’s paired-t test]. **I**. Voltage clamp recordings from representative WT (top) and YAC128 (bottom) CPNs at -70 mV in the presence of TTX (500 nM) and PTX (50 μM) (as before), with the addition of CPA (30 μM) and in the absence of extracellular Ca^2+^. Note in all such recording, CPNs were incubated in this CPA-containing, 0 mM Ca^2+^ ECSF for a minimum of 10 min. **J**. In CPA (30 μM) and in the absence of extracellular Ca^2+^, mEPSCs were significantly less frequent in YAC128 CPNs [3.3 ± 0.6 Hz (n=14; 4 cultures)] compared WT CPNs [8.4 ± 1.5 Hz (n=17; 5 cultures)] under like conditions [t(29)=2.952; p=0.0062; Student’s unpaired-t test]. **K**. In CPA (30 μM) and in the absence of extracellular Ca^2+^, YAC128 CPNs showed a trend towards a lower mean mEPSC amplitude than in WT (WT: 26.7 ± 5.2 pA, YAC128: 16.1 ± 3.3 pA), but this did not reach statistical significance [t(27)=1.706; p=0.0995; Student’s unpaired-t test].

### Removing extracellular Ca^2+^ and blocking the ER SERCA pump substantially reduces the mEPSC frequency in YAC128 but not WT cultures

If an ongoing release of presynaptic ER Ca^2+^ in YAC128 cultures occludes potentiation of synaptic glutamate release by low dose ryanodine and caffeine, blocking ER Ca^2+^ release should reduce synaptic glutamate release in YAC128 cultures. To test this, we pre-incubated cultures with extracellular fluid (ECF) containing the sarco/endoplasmic reticulum Ca^2+^-ATPase (SERCA) pump inhibitor cyclopiazonic acid (CPA) (30 μM) to deplete ER Ca^2+^ stores. Since ER Ca^2+^ depletion can increase cytosolic Ca^2+^ by engaging the store operated Ca^2+^ response, and this effect can increase mini neurotransmitter release (Emptage et al., 2001), we performed these experiments in the absence of extracellular Ca^2+^. Under these conditions, YAC128 CPNs showed a mean mEPSC frequency of 3.33 ± 0.57 Hz (n=14), significantly lower than seen in WT CPNs under identical conditions 8.41 ± 1.50 Hz (n=17) (Student’s unpaired t-test; p=0.0062) **(Figure 2I-K)**. These results suggest that an ER calcium leak into the cytoplasm elevates mini vesicular glutamate release from YAC128 CPN terminals, and that in the absence of this effect, the intrinsic vesicular release probability is actually reduced in YAC128 compared to WT cortical terminals.

### Presynaptic Ca^2+^ sparks and waves are more frequent in YAC128 cultures

To directly monitor presynaptic Ca^2+^ dynamics, we next fused GCaMP6-M (T.-W. Chen et al., 2013) with the rat synaptophysin protein via a small glycine-serine linker to generate a genetically-encoded Ca^2+^ sensor that preferentially localizes to presynaptic terminals. We expressed this rat synaptophysin-tagged GCaMP6-M construct (rSyph-GCaMP6m) in neurons in cortical cultures and performed Ca^2+^ imaging experiments, first in the presence of TTX (500 nM) to relate presynaptic Ca^2+^ signaling to our mEPSC findings (above). Remarkably, both YAC128 and WT cultures showed spontaneous axonal Ca^2+^ sparks, often beginning in single boutons, then initiating slow Ca^2+^ waves spreading to neighboring boutons **(Figure 3A, B)**. In some cases, the above waves slowly traversed axons encompassing entire 63X (178.6 μm x 113.1 μm) imaging fields over multiple seconds of imaging. Conversely, axonal Ca^2+^ events in the absence of TTX could not be temporally resolved (owing to relatively slow GCaMP kinetics) and appeared simultaneously across all the boutons of a given imaged axon. These mini presynaptic Ca^2+^ events in TTX were also strikingly long lasting at individual boutons - on average 5 - 10 times longer than that of typical events seen in the absence of TTX **(Figure 3C, D)**. These miniature presynaptic Ca^2+^ events in TTX were more than three times as frequent in YAC128 cultures than in WT cultures **(Figure 3E)**, but event DF/F amplitudes did not significantly differ between genotypes **(Figure 3F)**. The detection algorithm used here (see methods) considered Ca^2+^ waves involving multiple axonal boutons as single events (as in **Figure 3A**). However, the mean event area was similar in both genotypes (not illustrated), suggesting similar numbers of axonal boutons were recruited on average by such events in YAC128 and WT cultures. In subsets of WT and YAC128 cultures we also expressed an mCherry-tagged PSD95 construct. In these cultures, we repeatedly observed clear colocalizations between rSyph-GCaMP6m-labeled boutons, spontaneously active in TTX, and mCherry-labeled dendritic spines (not illustrated). Thus although we cannot be certain that all boutons showing spontaneous Ca^2+^ events in TTX participate in synaptic connections, it is clear that such events are not restricted to ectopic boutons.

**Fig 3:**
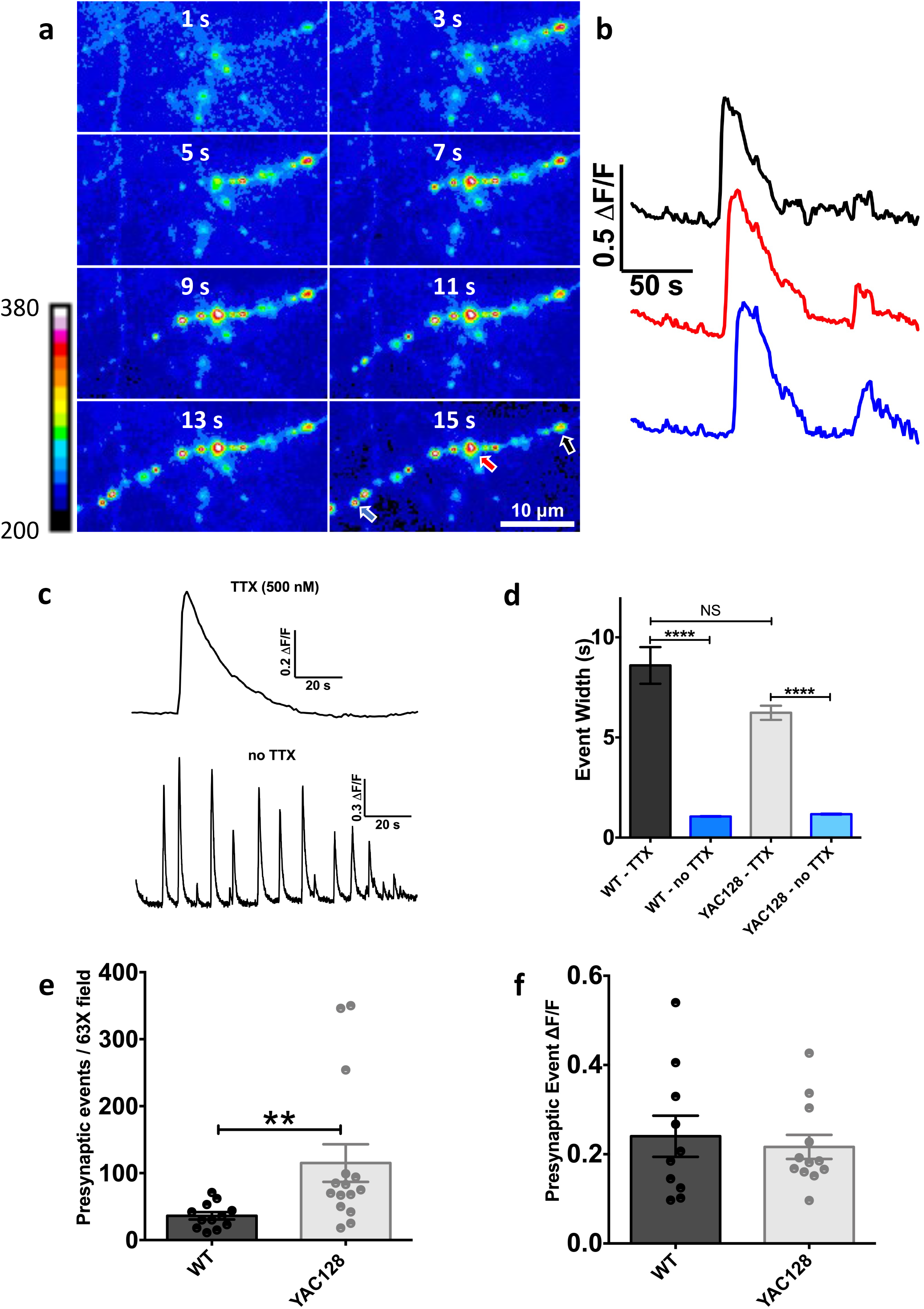
Axonal Ca^2+^ waves are more frequent in YAC128 Cortical Cultures. **A**. Portion of a YAC128 rat synaptophysin-tagged GCaMP6-M (rSyph-GCaMP6m)-expressing axon during a spontaneous Ca^2+^ wave in the presence of ECF containing TTX (500 nM). Images reflect raw GCaMP fluorescence values (arbitrary units) and comprise 15 s of a 4 min video. For illustrative purposes, images were binned temporally from the original 10 Hz movie, such that each 2 s frame reflects the average fluorescence of 20 successively acquired frames. **B**. Time course of DF/F (change in fluorescence over basal fluorescence) values extracted from 3 of the YAC128 rSyph-GCaMP6m-expressing axonal boutons (shown in panel **A** with color-coded arrows, panel labelled 15s). Signals were spatially averaged across elliptical regions of interest (ROI)s encompassing the indicated boutons and represent the entire 4 min imaging session. Note the multi-second delay in the propagation of signals between boutons and the long time course of individual events. **C**. Representative 3 min DF/F time course from a WT rSyph-GCaMP6m-expressing axonal bouton in TTX (500 nM) (top). Note the substantially longer duration of this event compared to representative events recorded without TTX from a separate WT rSyph-GCaMP6m-expressing bouton (bottom). **D**. Average half amplitude width of spontaneous Ca^2+^ events occurring in rSyph-GCaMP6m-expressing boutons in WT and YAC128 cortical cultures in the presence and absence of TTX (500 nM). Select comparisons were made with the Mann Whitney test because event widths in all groups failed the D’Agostino-Pearson omibus normality test. WT events were significantly longer with TTX [8.60 ± 0.92 s (n=164 events; 85 boutons; 4 cultures)] than without [1.06 ± 0.97 s (n=3725 events; 175 boutons; 4 cultures)] [p<0.0001 (approximate); Mann Whitney test]. Likewise, YAC128 half widths were significantly longer with TTX [6.23 ± 0.35 s (n=440 events; 153 boutons; 4 cultures)] than without [1.17 ± 1.46 s (n=5159 events; 422 boutons; 6 cultures)] [p<0.0001 (approximate); Mann Whitney test]. **E**. Numbers of detected spontaneous axonal events (as in Panels **A** and **B**) during 3 min recordings of 63 x objective (178.6 μm x 113.1 μm) fields imaged in the presence of TTX (500 nM). Action potential-independent events were significantly more frequent in YAC128 cortical cultures [115.9 ± 28.1 events/3 min (n=15 fields; 4 cultures)], compared to WT [36.3 ± 5.5 events/3 min (n=12 fields; 4 cultures)], [p=0.0016 (exact); Mann Whitney test]. **F**. Average peak DF/F amplitudes of all the spontaneous, action potential-independent axonal Ca^2+^ events imaged in a given 63 x objective field over 3 min from WT and YAC128 cortical cultures. Average event amplitudes were not significantly different between WT [0.240 ± 0.046 DF/F (n=10 fields; 4 cultures)] and YAC128 cultures [0.216 ± 0.027 DF/F (n=12 fields; 4 cultures)], [t(20)=0.465; p=0.647; Student’s unpaired-t test].

### Basal cytosolic Ca^2+^ is higher in YAC128 presynaptic boutons

In a subset of TTX experiments, we applied the Ca^2+^ ionophore ionomycin (10 μM) to cultures following GCaMP imaging. By equilibrating cytosolic Ca^2+^ to extracellular levels, this approach allowed quantification of GCaMP fluorescence in the presence of the known extracellular Ca^2+^ concentration (here 2 mM). Thus, smaller ionomycin-mediated increases in GCaMP fluorescence indicate relatively higher basal (pre-ionomycin) cytosolic Ca^2+^ concentrations (Lindhout et al., 2019). In both genotypes, ionomycin-mediated changes in presynaptic GCaMP fluorescence (DF/F) were significantly smaller in boutons that had shown at least one spontaneous Ca^2+^ event in the previous 3 min recording **(Figure 4A, D)**, indicating higher basal cytosolic Ca^2+^ concentrations in these spontaneously active boutons. Furthermore, overall ionomycin responses were significantly smaller in YAC128 cultures than in WT cultures when comparing the entire population of boutons imaged in a culture (both spontaneously active and inactive) **(Figure 4E)**.

**Fig 4:**
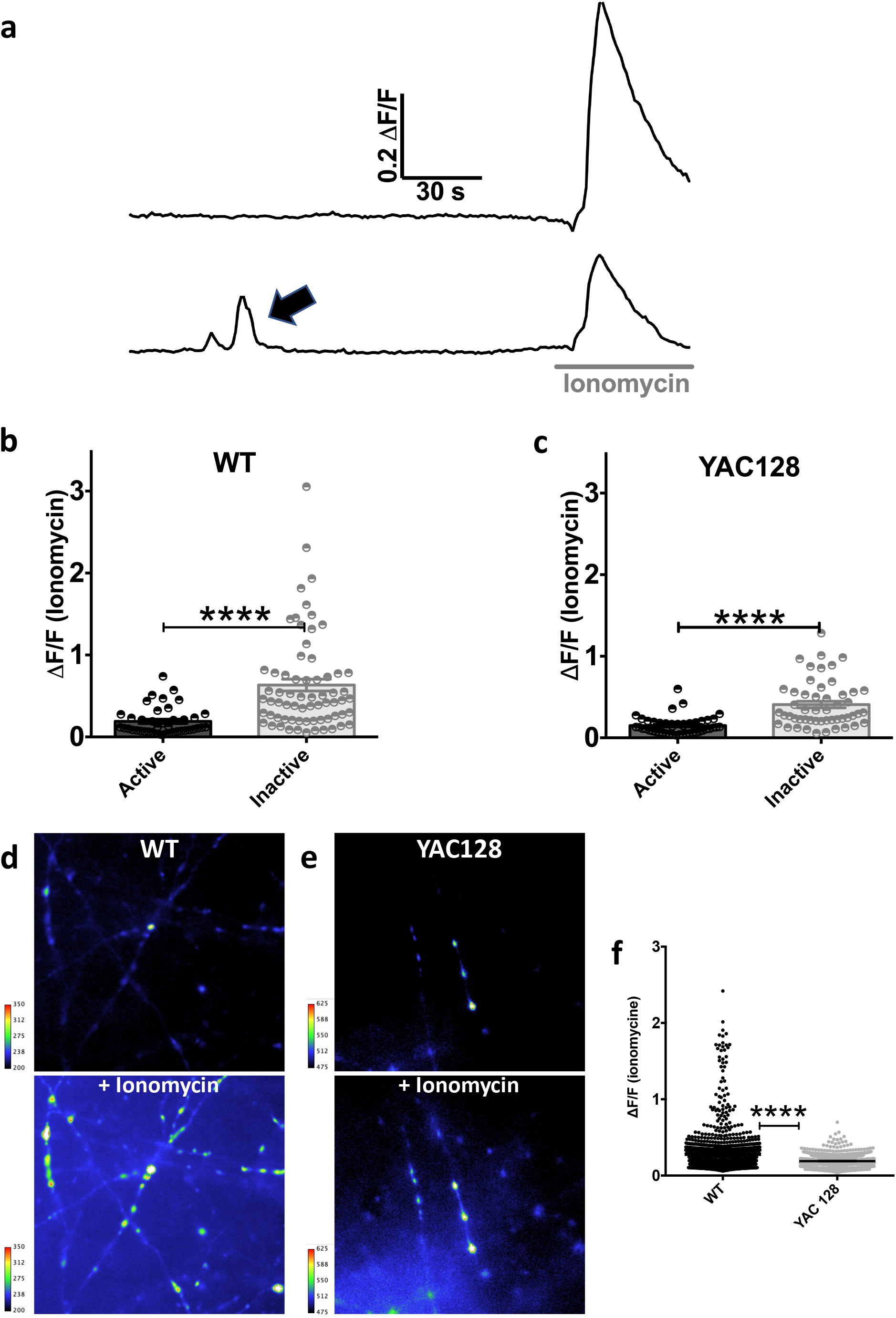
Boutons spontaneously active in TTX show relatively elevated cytosolic Ca^2+^ in both genotypes. Resting cytosolic Ca^2+^ concentrations are higher in the overall YAC128 bouton population. **A**. DF/F time courses from two rSyph-GCaMP6m-expressing axonal boutons from a representative YAC128 culture in the presence of TTX (500 nM). The first bouton (top trace) is inactive during the 3 min imaging session, while the second bouton (bottom trace) shows a clear spontaneous event (black arrow). Subsequent application of the Ca^2+^ ionophore ionomycin (10 μM) elicits a smaller Ca^2+^ response in the spontaneously active bouton, suggesting a higher basal cytosolic Ca^2+^ concentration (pre-ionomycin) in the active bouton. **B**,**C:** Ionomycin responses of rSyph-GCaMP6m-expressing boutons in the presence of TTX following a 3 min imaging session in WT (**B**) and YAC128 (**C**) cultures. Boutons were classified as active or inactive based on the presence, or lack thereof, of at least one spontaneous Ca^2+^ event during the initial 3 minute recording. Ionomycin responses were significantly larger in the population of inactive boutons [WT: 0.633 ± 0.072 DF/F (n=67 boutons); YAC128: 0.410 ± 0.040 DF/F (n=51 boutons)], vs active boutons [WT: 0.191 ± 0.026 DF/F (n=41 boutons; 4 cultures), p<0.0001 (exact) by Mann Whitney test; YAC128: 0.149 ± 0.016 DF/F (n=46 boutons), p<0.0001 (exact) by Mann Whitney test]. 4 cultures for WT and 4 cultures for YAC128. **D, E:** Responses of rSyph-GCaMP6m-expressing axons from a representative WT (**D**) and YAC128 (**E**) cortical culture to ionomycin (10 μM). Images are from a 4 min imaging session, with ionomycin application at 3 min and reflect raw GCaMP fluorescence values (arbitrary units) (as above). To minimize impacts of photobleaching, the top (before) image is the frame immediately prior ionomycin. The bottom image is a maximum projection of all frames acquired 30s prior and following ionomycin treatment and reflect each pixel’s maximum ionomycin response. The visible grey-value dynamic range is the same in panels **D** and **E**, (150 increments of the total unprocessed 14 bit-depth), however the 0 value in **E** has been adjusted so that brightness of the before image is comparable to that of its WT counterpart in **D**; this is so the reader can better appreciate the YAC128 culture’s smaller relative change in fluorescence. **F**. Ionomycin responses of the all WT and YAC128 rSyph-GCaMP6m-expressing axons automatically detected with the aQua software suite (see methods for reference). The WT bouton population showed a significantly greater ionomycin-mediated change in fluorescence [0.272 ± 0.006 DF/F (n=1681 boutons; 4 cultures)] compared to the YAC128 population [0.190 ± 0.002 DF/F (n=1462 boutons; 4 cultures)], [p<0.0001 (exact); Mann Whitney test].

### Caffeine increases basal Ca^2+^ and miniature events in WT, but not YAC128 cortical boutons

The slow kinetics of the above miniature presynaptic Ca^2+^ events are consistent with ER-mediated Ca^2+^ waves reported in postsynaptic neuronal compartments (Ross, 2012). We next tested whether these presynaptic Ca^2+^ events were affected by caffeine (1 mM), an ER ryanodine receptor agonist that substantially increased mEPSC frequencies in WT, but not YAC128 cultures (above). In the presence of TTX, caffeine (1 mM) significantly increased the presynaptic Ca^2+^ event frequency in WT cultures **(Figure 5A)**, but did not significantly alter event frequency in YAC128 cultures **(Figure 5B)**.

**Fig 5:**
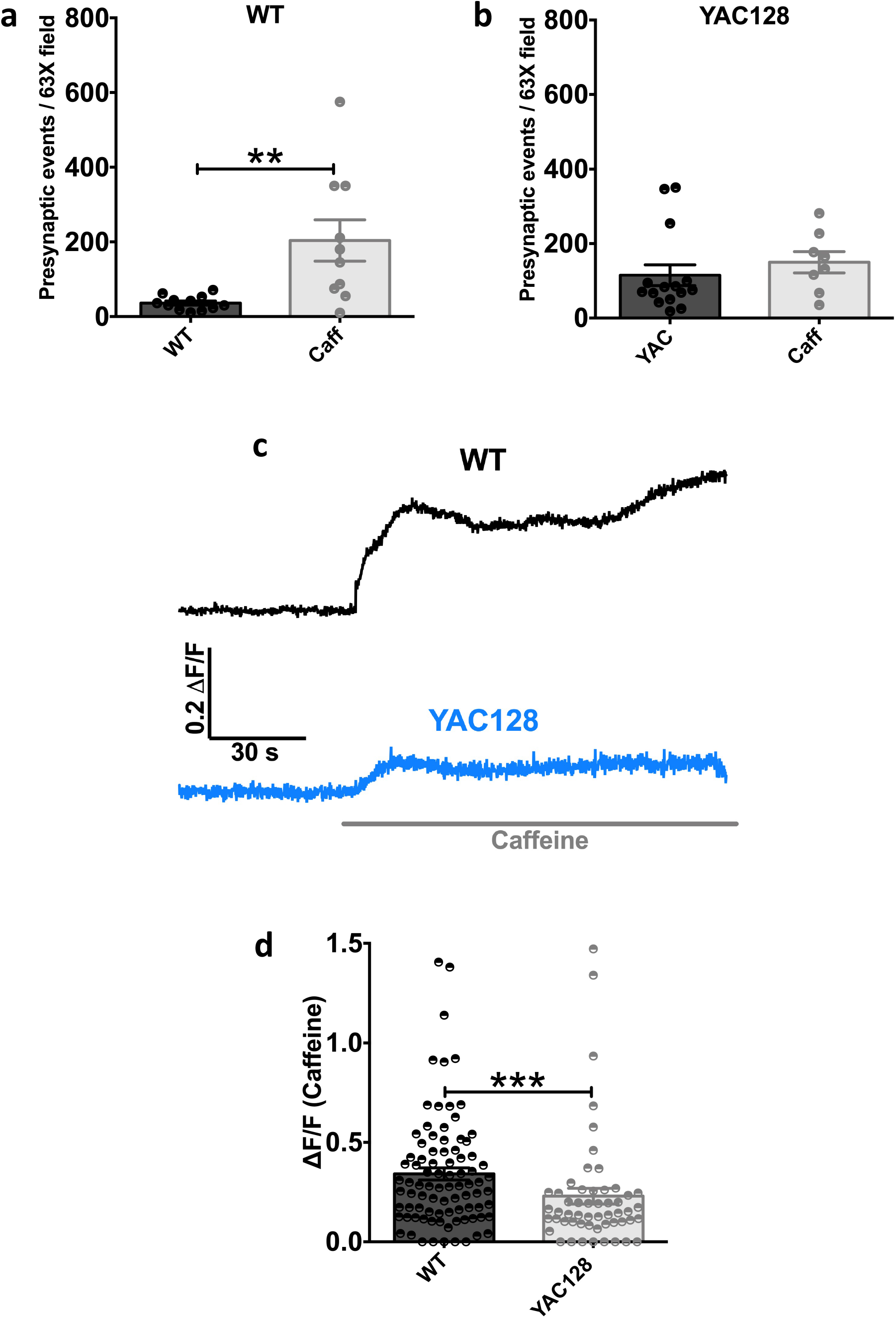
Caffeine increases the frequency of spontaneous Ca^2+^ events and the resting Ca^2+^ concentration in WT, but not YAC128, cortical axonal boutons. **A, B:** Numbers of detected spontaneous rSyph-GCaMP6m axonal events in the presence TTX (500 nM) during 3 min recordings of 63 x objective fields of WT (**A**) and YAC128 (**B**) cortical cultures (as above). The control group consists of the same culture fields shown in **Figure 6: E**, and is compared to separate WT (**A**) or YAC128 (**B**) culture fields imaged in the presence of caffeine (1 mM) under otherwise like conditions. WT axons showed significantly more action potential-independent events in the presence of caffeine, than in its absence [203.8 ± 55.2 events/3 min (n=10 fields; 5 cultures)] vs [36.3 ± 5.5 events/3 min (n=12 fields; 4 cultures)] respectively [t(20)=3.316; p=0.0034; Student’s unpaired-t test]. Numbers of detected YAC128 axonal events were not significantly different in the presence or absence of caffeine [115.1 ± 28.21 events/3 min (n=15 fields; 4 cultures)] vs [150.0 ± 28.55 events/3 min (n=8 fields; 4 cultures)] respectively [p=0.2188 (exact); Mann Whitney test]. **C**. DF/F time courses from representative WT (top) and YAC128 (bottom) rSyph-GCaMP6m-expressing axonal boutons in the presence of TTX (500 nM) before and immediately following application of caffeine (1 mM). For this illustration, boutons were selected which did not show spontaneous events. Rather, note the gradual sustained increase in rSyph-GCaMP6m fluorescence, which is of larger magnitude in the WT bouton. **D**. Caffeine responses of WT and YAC128 rSyph-GCaMP6m-expressing boutons in the presence of TTX as illustrated in panel **C**. Boutons showing spontaneous events **(as in Figure 6)** were excluded as the long time course of such events complicated quantification of steady-state increases in DF/F. Caffeine application elicited a significantly greater increase in rSyph-GCaMP6m fluorescence in WT boutons [0.342 ± 0.030 DF/F (n=87 boutons)] compared to YAC128 [0.229 ± 0.039 DF/F (n=54 boutons)] [p=0.0004 (exact); Mann Whitney test]. For A, B and D: 5 cultures for WT and 5 cultures for YAC128.

Significant photobleaching of the rSyph-GCaMP6m construct occurred with prolonged imaging; this combined with the relative infrequency of mini presynaptic Ca^2+^ events necessitated imaging of separate fields for the above control and caffeine comparisons. We next performed within-bouton measurements of basal rSyph-GCaMP6m fluorescence before and immediately following application of caffeine; the relatively short duration of these experiments made before and after measurements feasible. A clear slow increase in basal rSyph-GCaMP6m fluorescence was seen in most WT boutons following caffeine (1 mM) application **(Figure 5C)**, which was generally absent or reduced in YAC128 boutons **(Figure 5D)**. Overall a significantly greater caffeine (1 mM)-mediated increase in rSyph-GCaMP6m fluorescence was seen in WT boutons compared to YAC128 **(Figure 5E)**.

### Action potential-dependent Ca^2+^ signals in presynaptic cortical terminals Are less frequent in YAC128 cultures

We next used the rSyph-GCaMP6m construct in the absence of TTX to examine presynaptic action potential-dependent Ca^2+^ signals in WT and YAC128 cortical cultures. When neuronal action potential firing was intact, rSyph-GCaMP6m -expressing boutons in both WT and YAC128 cultures were dominated by presumably voltage-gated Ca^2+^ channel-mediated signals **(Figure 6A, B)**. These signals were more frequent than those seen in the presence of TTX, but of far shorter duration. Interestingly, these activity-dependent Ca^2+^ events were nearly twice as frequent in WT axonal boutons, compared to those in YAC128 cultures **(Figure 6A, C and E)**. Ryanodine (5 μM) modestly but significantly reduced the frequency of these events in WT cultures (by 17 %) **(Figure 6A, B and E)**. Conversely, 5 μM ryanodine elicited a modest, but significant increase in the frequency of these events in YAC128 cultures **(Figure 6C-E)**.

**Fig 6:**
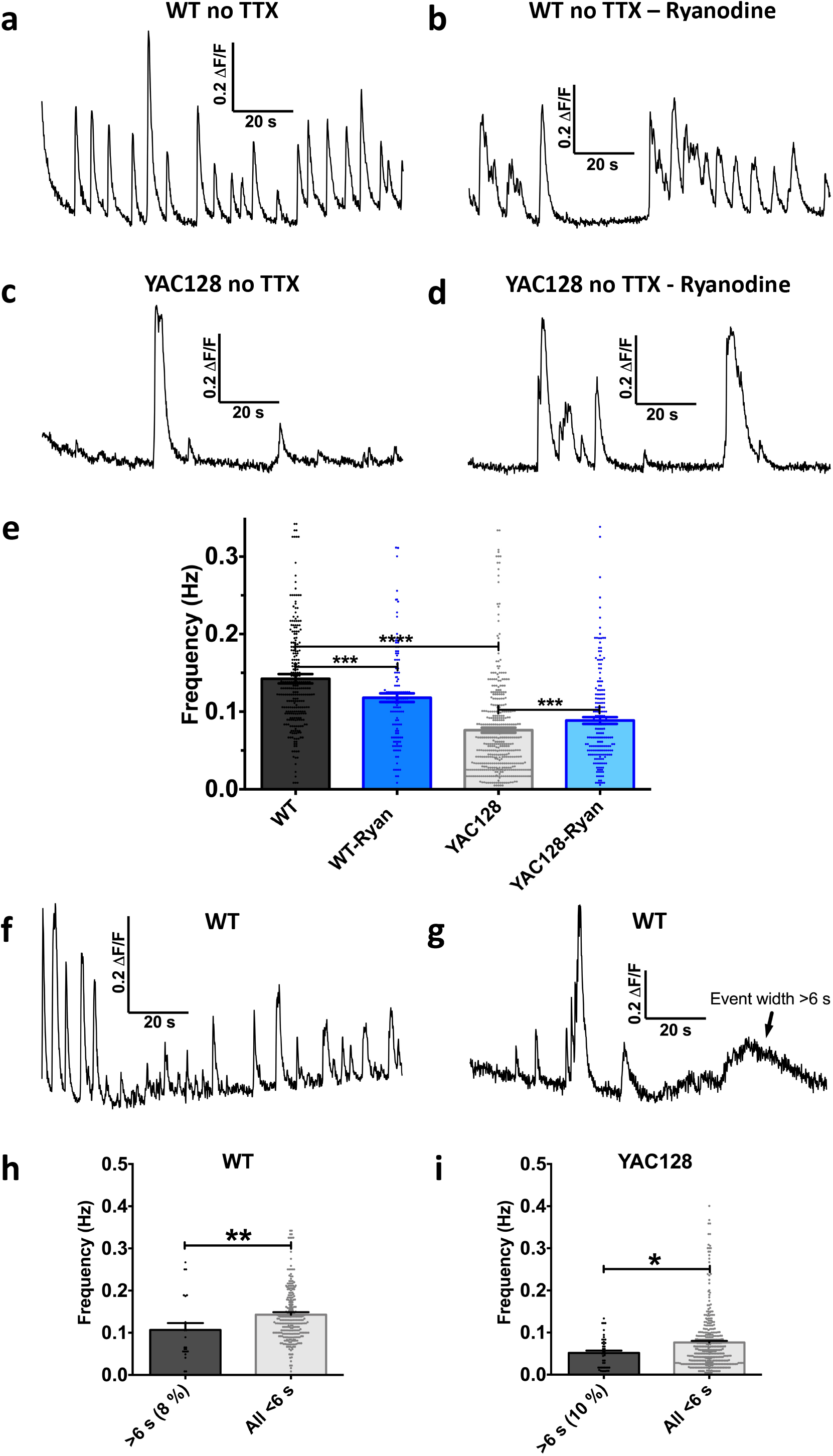
In the absence of TTX, presynaptic calcium events are more frequent in WT, compared to YAC128, cultures. **A, B:** DF/F time course from representative WT rSyph-GCaMP6m-expressing axonal boutons in the absence of TTX, under control conditions (A) and in presences of 5 μM ryanodine (B). Note these are not the same boutons, as experiments illustrated in this figure were not paired. **C, D:** DF/F time course from two different representative YAC128 rSyph-GCaMP6m-expressing axonal boutons in the absence of TTX, one under control conditions (C) and the other in the presence of 5 μM ryanodine. (D) **E:** Mean Ca^2+^ event frequencies of rSyph-GCaMP6m-expressing boutons in the absence of TTX in WT and YAC128 cortical cultures in the presence and absence of ryanodine (5 μM). Select comparisons were made with the Mann Whitney test because event widths in all groups failed the D’Agostino-Pearson omibus normality test. Events were significantly more frequent in WT boutons [0.1424 ± 0.0061 Hz (n=272 boutons; 5 cultures)] than in YAC128 boutons [0.0761 ± 0.0044 Hz (n=423 boutons; 6 cultures)] [p<0.0001 (approximate); Mann Whitney test]. Events were significantly less frequent in WT boutons with ryanodine (5 μM) [0.1179 ± 0.0033 Hz (n=120 boutons, 3 cultures)] than without (above) [p=0.0038 (approximate); Mann Whitney test]. Conversely, event frequencies were significantly more frequent in YAC128 boutons with ryanodine (5 μM) [0.0886 ± 0.0044 Hz (n=189 boutons, 4 cultures)] than without (above) [p=0.0003 (approximate); Mann Whitney test]. **F, G:** DF/F time course from two representative rSyph-GCaMP6m-expressing axonal boutons from the same WT culture field in the absence of TTX. Bouton (F) shows only faster events (lasting < 6 s), while in bouton (G) a clear slow event (lasting >6 s) is present. Note the substantially lower overall event frequency in (G). **H, I:** Ca^2+^ event frequencies of WT (H) and YAC128 (I) rSyph-GCaMP6m-expressing boutons (as above), grouped based on the presence or absence of one or more slow events (> 6 s at half peak amp). (H): 23/292 (7.9 %) WT boutons showed at least on such slow event with an overall mean bouton frequency of 0.107 ± 0.016 Hz (n=23), significantly lower than WT boutons lacking slower events [0.143 ± 0.006 Hz (n=269)] [p=0.0058 (exact); Mann Whitney test] (n=292 boutons, 5 cultures). (I) 44/423 (10.4 %) YAC128 boutons showed at least one such slow event with an overall mean bouton frequency of 0.052 ± 0.005 Hz (n=44), significantly lower than YAC128 boutons lacking slower events [0.076 ± 0.004 Hz (n=379)] [p=0.0058 (exact); Mann Whitney test] (n=423 boutons, 6 cultures).

As discussed above, rSyph-GCaMP6m events in the presence of TTX were typically much longer in duration than action potential-dependent events in both genotypes and appeared to be restricted to a subset of boutons with higher resting Ca^2+^ concentrations. Interestingly, similar strikingly long-lasting events continued to occur in subsets of boutons in the absence of TTX. In an attempt to identify boutons exhibiting these slow Ca^2+^ events, we next categorized boutons imaged in the absence of TTX based on whether one or more Ca^2+^ events with a duration greater than 6 s (measured at half peak amplitude) occurred during a 3 min recording. 6 s was chosen as the cutoff value, because it was more than 5 standard deviations greater than the mean event duration seen in WT and YAC128 boutons in the absence of TTX, but comparable to average event durations in TTX in both genotypes: [8.60 ± 0.92 s (n=164) and 6.23 ± 0.35 s (n=440) in WT and YAC128 boutons respectively]. Without TTX, 7.9 % (23/292) of WT boutons showed one or more events lasting longer than 6 s, while under identical conditions 10.4 % (44/423) of YAC128 boutons showed at least one such event. Interestingly in both genotypes, overall event frequencies were significantly lower in boutons showing one or more slow events (>6 s) than in boutons showing only short duration events **(Figure 6F-I)**. Assuming boutons displaying these long lasting Ca^2+^ events correspond to the same population that is spontaneously active in TTX, these results suggest action potential-dependent Ca^2+^ events are reduced in the bouton population with higher resting cytosolic Ca^2+^ levels and spontaneous ER release events. Taken together, experiments up to this point suggest that releasing presynaptic ER Ca^2+^ elevates miniature vesicular glutamate release, but can also reduce activity-dependent presynaptic Ca^2+^ influx, and imply that the spontaneous or pharmacological release of presynaptic ER Ca^2+^ oppositely regulates miniature and action potential-dependent glutamate release.

### Low dose ryanodine reduces evoked glutamate release in WT-but not YAC128-derived brain slices

We next performed experiments in cortical-striatal *ex-vivo* brain slices prepared from 2 – 3 month old WT and YAC128 mice expressing the fluorescent glutamate sensor iGluSnFR in striatal neurons (Parsons et al. 2016; Koch et al. 2018). This preparation allowed direct optical measurement of glutamate release in the striatum, independent of postsynaptic neuronal properties. Striatal iGluSnFR signals evoked by stimulating cortical axons of the corpus callosum were significantly decreased by ryanodine (5 μM) in WT, but not YAC128 slices **(Figure 7)**. These data are consistent with results in culture (Figs. 6), suggesting that AP-dependent glutamate release is reduced in YAC128 as a consequence of a tonic Ca^2+^ leak from ER stores

**Fig 7:**
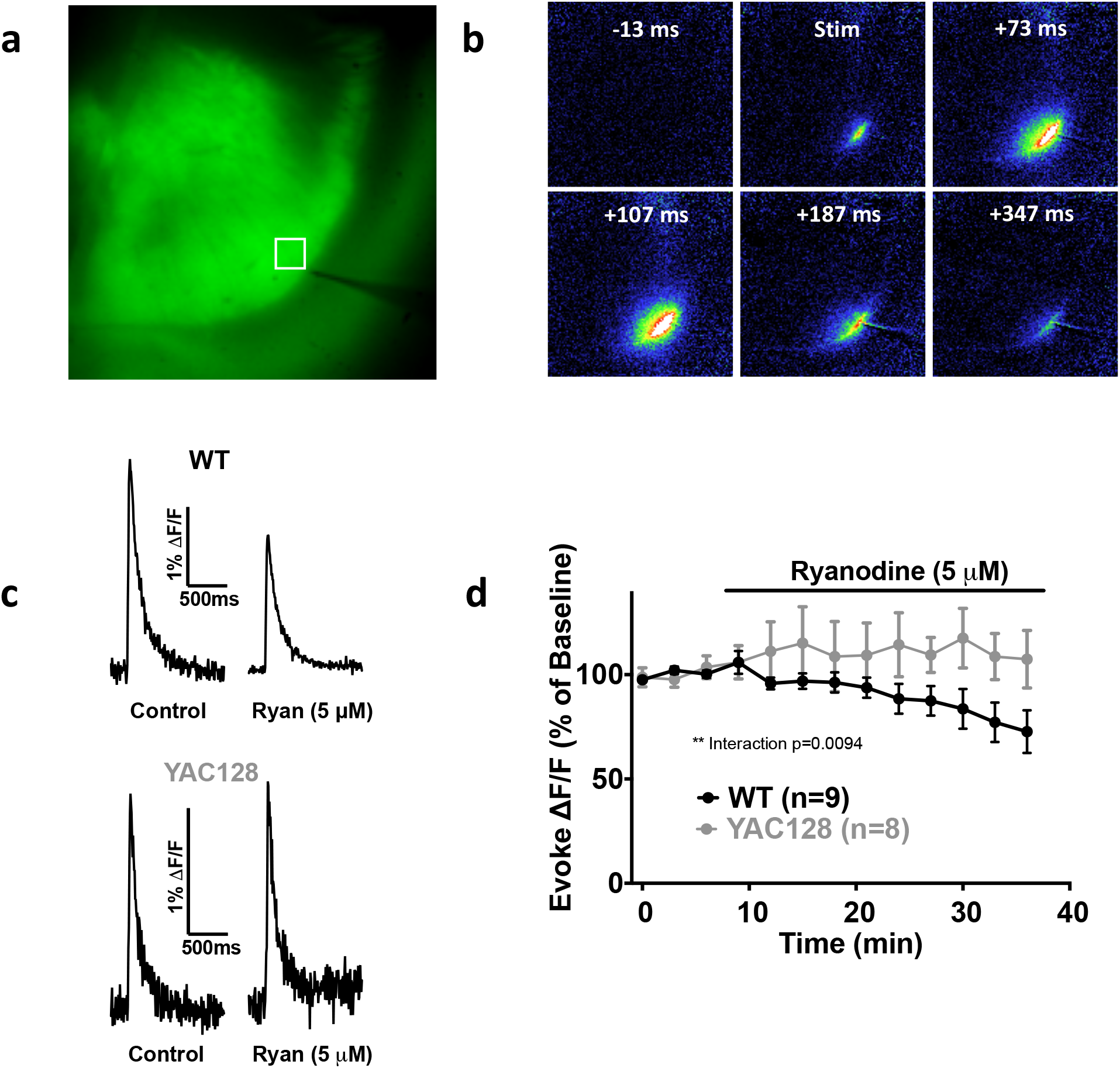
Ryanodine reduces evoked striatal iGluSnR responses in WT, but not YAC128, brain slices. **A**. Image of raw iGluSnFR fluorescence from a representative WT mouse brain slice with viral-mediated expression of iGluSnFR in its dorsal striatum. Note the square ROI over which evoked spatially averaged DF/F signals were measured, and the tungsten monopolar stimulating electrode placed in the corpus callosum. **B**. A DF/F image montage from the WT brain slice (shown in A), illustrating a striatum iGluSnFR response evoked by electrically stimulating cortical axons in the adjacent corpus callosum. **C**. Striatum evoked DF/F iGluSnFR responses from a representative WT (top) and YAC128 (bottom) brain slice before and in the presence of ryanodine (5 μM). Ryanodine elicits a substantial reduction in the peak WT, but not YAC128, iGluSnFR response. **D**. Peak evoked striatum iGluSnFR responses from WT and YAC128 slices before and during treatment of slices with ryanodine (5 μM). Responses were measured repeatedly at 3 min intervals and were normalized to the average of each slice’s 3 baseline (before drug) measurements. Ryanodine application decreased the amplitude of the evoked iGluSnFR response in WT (n=9 slices; 9 mice), but not YAC128 (n=8 slices; 8 mice). The effect of ryanodine on WT slices was significantly different than in YAC128 slices, based on a significant time-genotype interaction (p=0.0094, repeated measure (time) two way ANOVA).

## Discussion

Presynaptic neurotransmitter release and consequent postsynaptic signaling largely underlies communication between neurons, thus forming the basis of brain circuitry. Neurotransmitter release in turn, can be broadly divided into action potential-dependent or independent forms. The former requires presynaptic voltage-gated Ca^2+^ channel activation coordinated by sodium action potentials; while the latter persists in the absence of neuronal activity. Although miniature neurotransmission is poorly understood compared to its activity-dependent counterpart, it is increasingly accepted to subserve clear physiological functions (Frank, Kennedy, Goold, Marek, & Davis, 2006; McKinney, Capogna, Dürr, Gähwiler, & Thompson, 1999; Sutton et al., 2006). Moreover, miniature release can be regulated relatively independently of action potential-dependent release and may even be mediated by distinct vesicular pools (Fredj & Burrone, 2009; Sara, Virmani, Deák, Liu, & Kavalali, 2005).

Alterations in synaptic signaling processes, particularly relating to the excitatory transmitter glutamate, have been repeatedly reported in models of HD (Raymond et al., 2011; Sepers et al., 2017; Tyebji & Hannan, 2017) and other neurodegenerative diseases (R. Wang & Reddy, 2017). Increased cell-surface expression of extra-synaptic NMDA receptors by SPNs favors excitotoxic postsynaptic glutamate-mediated signaling in HD models (Milnerwood et al., 2010). Mounting evidence also suggests glutamate release from cortical afferents is altered in HD; however, the direction of this alteration appears to be disease-stage-dependent (Cepeda et al., 2003; Joshi et al., 2009).

The endoplasmic reticulum (ER), a continuous intracellular membrane-system involved in Ca^2+^ storage and protein synthesis, is expressed in all neuronal processes including axons and presynaptic boutons. Increased cytosolic Ca^2+^-release from the ER has been shown in HD models, as a result of a mutant huntingtin protein (mHTT)-mediated increase in IP3 receptor responsiveness (Tang et al., 2003) and a constituent ryanodine receptor Ca^2+^ leak (Suzuki et al., 2012). Previous HD studies have focused on the postsynaptic ER, however Ca^2+^ release from the presynaptic ER is known to modulate neurotransmitter release (Emptage et al., 2001; Llano et al., 2000). Results of our study suggest mHTT expression alters ER Ca^2+^ handling at stores in close proximity to cortical presynaptic terminals, thereby altering glutamate release. This mechanism clearly increased the frequency of Ca^2+^-dependent, action potential-independent (mini) synaptic glutamate events in our YAC128 model. Conversely, our findings suggest reduced action potential-dependent glutamate release from YAC128 cortical terminals. As we have focused exclusively on glutamatergic synapses, future experiments will be required to determine if this mechanism similarly affects the release of other neurotransmitters and neuromodulators.

### Spontaneous axonal Ca^2+^ signaling increases the YAC128 miniature glutamate release frequency

CPNs in YAC128 cortical cultures at DIV14 and DIV18 showed higher mEPSCs frequencies compared to CPNs in age-matched WT cultures. However by DIV 21, mEPSC frequencies were similar between WT and YAC128 CPNs. This pattern is reminiscent of SPN mEPSC frequencies in YAC128 cortical-striatal co-cultures, which were also higher relative to WT at early DIV ages, but comparable to WT by DIV21 (Buren et al., 2016). In the co-culture model, the total dendritic length of YAC128 SPNs was reduced relative to WT at DIV21, indicating reduced total SPN synapse numbers may have masked ongoing elevated glutamate release rates from individual YAC128 cortical terminals. We suspect that in our cortical cultures, degenerative changes at DIV21 likewise obscured the elevated YAC128 CPN mEPSC frequencies clearly seen at earlier DIV ages. Notably in DIV18-aged cortical cultures, the time point at which we performed mechanistic experiments, no genotype differences in CPN synapse density and dendritic morphology were evident, indicating that differences in mEPSC frequencies likely reflected altered rates of release at individual presynaptic sites. In any case, increased rates of miniature glutamate release appear to be an early and perhaps enduring phenotype of cultured YAC128 CPNs.

Caffeine or low dose ryanodine, which release ER Ca^2+^ by opening ryanodine receptors, failed to increase the mEPSC frequency in YAC128 CPNs, despite robustly increasing the WT mEPSC frequency. This result, in concert with the increased basal YAC128 mEPSC frequency, suggests an ongoing spontaneous Ca^2+^ store release in YAC128 cultures occludes facilitation of the mEPSC frequency by drug-mediated Ca^2+^ store release. The substantially lower CPN mEPSC frequency seen in YAC128 cultures incubated with the ER SERCA pump inhibitor CPA in nominally zero extracellular Ca^2+^, is also consistent with a strong ER-derived Ca^2+^-dependence of YAC128 mini glutamate release. Conversely, mEPSC frequencies in WT cultures were not substantially reduced under identical conditions; this resulted in significantly higher WT than YAC128 mEPSC frequencies in CPA and 0 mM extracellular Ca^2+^, notably opposite the relationship seen without CPA in standard (2 mM) extracellular Ca^2+^. This was unexpected given overall WT and YAC128 CPN synapse numbers were similar in these cultures, but may indicate a reduction in functional YAC128 synapses, not apparent in fixed culture images, or deficits in the YAC128 synaptic release machinery, as suggested in previous studies (Morton & Edwardson, 2001; Morton, Faull, & Edwardson, 2001). In either case, enhanced Ca^2+^-dependent mini glutamate release in YAC128 cultures clearly surmounts any such deficits. These results also suggest mini glutamate release in WT cultures is largely Ca^2+^-independent. In contrast, Xu et. al (2009) reported mini glutamate release in similar cortical cultures persisted in the absence of extracellular Ca^2+^, but was nearly completely dependent on internal Ca^2+^stores and blocked by preincubation with the membrane-permeant Ca^2+^ chelator BAPTA-AM (Xu, Pang, Shin, & Südhof, 2009). If this were true in our cultures, we would expect Ca^2+^ store depletion (with CPA), in concert with removal of extracellular Ca^2+^, to similarly abolish mini release. However CPA is reported to only partially deplete presynaptic ER Ca^2+^ (de Juan-Sanz et al., 2017), therefore a role of residual ER Ca^2+^ in maintaining mini release in WT cultures under these conditions cannot be excluded.

Despite a clear role of ER Ca^2+^ in mini release in a variety of neuronal preparations, the temporal and spatial dynamics of action potential-independent presynaptic ER Ca^2+^ signals are largely unclear. Imaging the rSyph-GCaMP6m probe in TTX revealed spontaneous, presynaptic Ca^2+^ events that often initiated Ca^2+^ waves traversing cortical axons. These events were more frequent in YAC128 cultures and showed strikingly slow kinetics in both genotypes, consistent with ER Ca^2+^ waves reported in dendrites and non-neuronal cell-types (Ross, 2012). Caffeine substantially increased the mini axonal Ca^2+^ event frequency in WT cultures to levels comparable to that seen in YAC128 cultures, but failed to significantly change the event frequency in YAC128 cultures. The congruence of these results with our mEPSC findings (above) suggests these axonal Ca^2+^ events underlie the increased YAC128 mini glutamate release frequencies.

Basal cytosolic Ca^2+^ concentrations were higher in axonal boutons showing spontaneous Ca^2+^ activity in TTX, compared to inactive boutons, in both genotypes. Elevated cytosolic Ca^2+^ levels may therefore be required to precipitate the initial ryanodine or IP3 receptor opening needed to incite the regenerative ER Ca^2+^-induced Ca^2+^ release we presume underlie these axonal Ca^2+^ events. Interestingly, the restriction of spontaneous Ca^2+^ signals to a subset of cortical boutons with higher basal Ca^2+^ concentrations may mean mini glutamate release is not uniform across cortical boutons, but favored from this population of spontaneously active boutons. Indeed, subpopulations of synapses favoring mini release have been described in other neuronal systems (Atasoy et al., 2008; Peled, Newman, & Isacoff, 2014; Reese & Kavalali, 2016). We suspect the higher basal cytosolic Ca^2+^ concentrations in the overall YAC128 axonal bouton population meant a greater proportion were capable of sustaining spontaneous Ca^2+^ activity. Caffeine application increased the baseline cytosolic Ca^2+^ concentration and frequency of spontaneous events in WT, but not YAC128 boutons, consistent with this idea.

### Action potential-dependent glutamate release is suppressed due to presynaptic ER depletion

Regardless of genotype, striking differences were seen in the rSyph-GCaMP6m signals that predominated in the presence versus absence of TTX. Most rSyph-GCaMP6m-expressing boutons in the absence of TTX showed numerous spontaneous events during standard 3 min imaging sessions. However, only subsets of boutons were spontaneously active in TTX, with active boutons characterized by higher resting cytosolic Ca^2+^ concentrations. Furthermore, although Ca^2+^ events at individual boutons in TTX were less frequent, they lasted many times longer than corresponding action potential-dependent events. Interestingly in the absence of TTX, subsets of boutons of both genotypes showed longer-lasting Ca^2+^ events, like those that persisted in TTX, typically alongside faster action potential-dependent events. We suspect these boutons correspond to the population spontaneously active in TTX. Event frequencies in such boutons, in the absence of TTX, were significantly lower than in boutons lacking these longer-lasting events, further supporting the idea that subsets of cortical boutons favor either miniature or action potential-dependent glutamate release.

Although spontaneous axonal Ca^2+^ events were more common in YAC128 cultures in the presence of TTX, the opposite was true when experiments were performed without TTX, suggesting reduced activity-dependent glutamate release in YAC128 cultures. Reduced action potential-dependent axonal Ca^2+^ event frequencies in YAC128 cultures could reflect decreased CPN firing rates, altered transduction of sodium action potentials to bouton Ca^2+^ signals or some combination of these factors. 5 μM ryanodine significantly reduced the frequency of these action potential-dependent bouton Ca^2+^ events in WT cultures, though not to YAC128 levels, consistent with reduced action potential-dependent presynaptic Ca^2+^ signaling in YAC128 cultures being partly, but not entirely mediated, by spontaneous release of ER Ca^2+^. However, nuances of ryanodine’s pharmacology may have confounded this conclusion. Low dose ryanodine opens ryanodine receptors, but to a sub-maximal conductance state. Furthermore, ryanodine binding to a second low affinity receptor site mediates channel closure, an action that predominates at doses greater than 10 μM. Caffeine more effectively releases ER Ca^2+^, exemplified here in patch clamp mini EPSC experiments, and may therefore have been capable of lowering action potential-dependent event frequencies in WT to YAC128 levels. Unfortunately, caffeine was undesirable for these experiments because, unlike ryanodine, it is an antagonist at adenosine receptors, which are often expressed at presynaptic sites and inhibit VGCCs (Dunwiddie & Masino, 2001). We speculate that the ryanodine receptor population in YAC128 boutons favors the open conformation under basal conditions, consistent with the lack of caffeine responses in YAC128 boutons, and that under such conditions, ryanodine-mediated channel closure would be exaggerated. If so, this could account for the modest, but significant ryanodine-mediated increase in event frequencies in YAC128 boutons.

Precise mechanism(s) by which pharmacological or disease-mediated ER Ca^2+^ release might reduce activity-dependent presynaptic Ca^2+^ signaling remain uncertain. Indeed axonal or cell body ER stores could equally underlie this effect given the dependence of these signals on sodium action potentials. Dendritic ryanodine receptor activation has been shown to reduce neuronal firing rates via activation of a Ca^2+^-dependent K^+^ conductance (van de Vrede, Fossier, Baux, Joels, & Chameau, 2007). Alternatively, elevated cytosolic Ca^2+^ levels in YAC128 boutons (mediated by the release of axonal stores) might contribute to Ca^2+^-dependent VGCC inactivation, thereby reducing the coupling of cortical action potentials to presynaptic VGCC-mediated Ca^2+^ influx. Indeed, more generalized increases in cytosolic Ca^2+^ mediated by ER store release, have been shown to facilitate Ca^2+^-dependent inactivation of voltage-gated Ca^2+^ channels including the N and P/Q-types commonly expressed in presynaptic terminals and implicated in vesicular release (Budde, Meuth, & Pape, 2002; Cens, Rousset, Leyris, Fesquet, & Charnet, 2006). If this were the case in our system, the higher resting Ca^2+^ concentrations evident in YAC128 boutons would be expected to maintain local voltage-gated Ca^2+^ channels in a higher resting state of inactivation.

Low-dose ryanodine decreased the amplitude of striatum, glutamate-mediated iGluSnFR signals in WT brain slices, an effect absent or diminished in YAC128 slices, consistent with our findings in culture. There is a lack of consensus as to whether single action potentials can trigger Ca^2+^-induced Ca^2+^ release from the presynaptic ER (Emptage et al., 2001), or whether prolonged, repetitive action potential firing is required (de Juan-Sanz et al., 2017). It seems unlikely that the stimulation protocol used here appreciably contributed to the action potential-dependent discharge of the presynaptic ER, given the relatively low intensity stimulation used (repeated only twice at 100 Hz). Rather, we suspect a ryanodine-mediated increase in presynaptic Ca^2+^ decreased evoked glutamate release by inactivating presynaptic voltage-gated Ca^2+^ channels in WT slices, and that this process was occluded in YAC128 slices. However, future studies will be required to clarify mechanistic details of this process.

Taken together, the evidence presented here strongly suggests enhanced spontaneous release of presynaptic ER Ca^2+^ in YAC128 mice favors miniature glutamate release at the expense of evoked release.

### Implications for postsynaptic signaling

Mini glutamate release elicits postsynaptic NMDA receptor-mediated Ca^2+^ influx under physiologically relevant conditions (Beaulieu-Laroche & Harnett, 2018; Espinosa & Kavalali, 2009) and can mediate postsynaptic signaling distinct from that of action potential-dependent release (Sutton, Taylor, Ito, Pham, & Schuman, 2007). Indeed mini glutamate release may activate distinct populations of NMDAR receptors (Atasoy et al., 2008). In cultures, it was reported that miniature glutamate-mediated events become toxic to CPNs following prolonged silencing of neuronal activity with TTX (Fishbein & Segal, 2007). Future studies will be necessary to determine if and how the increased mini glutamate release shown here interacts with the well described alterations in postsynaptic NMDA receptor expression in the YAC128 and other HD-models, and whether a shift towards activity-independent glutamate release contributes to neurodegeneration in HD.

## Materials and Methods

### Culture preparation

All animal-related procedures were approved by and adhered to the guidelines of the University of British Columbia Committee on Animal Care and the Canadian Council on Animal Care (protocols A17-0295, A15-0069 and A19-0076). Cultures were prepared from both male and female embryonic day 17-18 pups from either wild-type (WT) FVB/N or transgenic yeast artificial chromosome-containing mice expressing the full-length human huntingtin genomic DNA with 128 CAG repeats (YAC128). YAC128 mice were maintained on the FVB/N background (homozygous line 55). Wildtype and YAC128 mice used for ex vivo slice experiments (below) and bred for culture preparation (above) were group housed under controlled conditions, free of know pathogens, at room temperature (22 – 24 °C), under a 12 hr light/dark cycle. Cortical cultures used in patch clamp electrophysiology and Ca^2+^-imaging experiments were prepared as previously described (Milnerwood et al., 2012; Smith-Dijak et al., 2019) and plated at a density of 225, 000 neurons/ml. In a subsets of experiments, a portion of the total 2.7 million cortical neurons (plated per 24-well culture) were transfected with transgenic reporters including: GFP; a synaptophysin-tagged GCaMP6-M construct (a generous gift from Dr. Anne Marie Craig, UBC); a postsynaptic density 95 (PSD-95)–tagged M-cherry construct; or a GFP-tagged internally expressed anti-PSD95 antibody (a generous gift from Dr. D.B. Arnold, University of Southern California; (Gross et al., 2013)).

### Electrophysiology

An Axopatch 200B amplifier and pClamp 9.2 software (Molecular Devices, Sunnyvale, CA) were used to acquire whole-cell patch clamp electrophysiology recordings. Data was digitized at 20 kHz and low-pass filtered at 1 kHz. For electrophysiology experiments, cultures were perfused with extracellular fluid (ECF) containing (in mM): 167 NaCl, 2.4 KCl, 10 glucose, 10 HEPES, 2 CaCl_2_ and 1 MgCl_2_; NaOH (1 mM) was used to adjust the pH to 7.30, and the osmolarity was adjusted to 305 – 310 mOsm. Tetrodotoxin (TTX) (500 nM) and picrotoxin (PTX) (50 mM) were added to this ECF to block sodium channel-mediated action potentials and GABA_A_ receptor-mediated currents respectively. Neurons were patched with borosilicate glass pipettes pulled to a tip resistance of 3-6 MΩ when back-filled with intracellular solution containing in (mM): 130 Cs-methanesulfonate, 5 CsCl, 4 NaCl, 1 MgCl, 10 HEPES, 5 EGTA, 5 QX-314 Cl, 0.5 Na-GTP, 10 Na-phosphocreatine, and 5 Mg-ATP (∼286 mOsm). During experiments, neurons were held at – 70 mV in voltage-clamp, with hyperpolarizing voltage steps (−10 mV) performed periodically to measure intrinsic membrane properties. Under these conditions AMPA receptor-mediated miniature excitatory postsynaptic currents (mEPSC)s appeared as transient inward current deflections. Recordings with a series resistance greater than 25 MΩ were excluded from analysis; typical values were between 15 – 20 MΩ. A minimum of 2 minutes following establishment of the whole cell configuration was allowed before experimental measurements, so neurons could fully dialyze with intracellular solution and achieve a stable membrane resistance and holding current. For experiments involving within-cell drug applications, a maximum 20 % change in series resistance between control and drug measurements was tolerated. In these cases, drugs were applied locally to the neuron with a fast perfusion system. Mini Analysis software (Synaptosoft) was used to detect mEPSCs and extract relevant parameters. A minimum of 100 and no more than 1000 mEPSC’s were analyzed per neuron per experimental condition.

### Cortical pyramidal neuron morphology

Cortical pyramidal neurons (CPN)s in WT and YAC128 cultures were labeled by transfecting a subset of neurons (1 of 2.7 million) with a cytoplasmic green fluorescent protein (GFP) (Addgene plasmid 37825) at the time of platting. Cultures were subsequently fixed at DIV17 - 19 and GFP-labeled CPN dendritic arbors imaged on a Zeiss Axiovert 200 M fluorescence microscope (20 x magnification, 0.8 NA), with a Zeiss 702 monochrome camera, using Zen software. Multiple image Z-stacks were acquired and X,Y-tiling was used to ensure entire dendritic arbors were visualized. Images were exported to Fiji-ImageJ for analysis by a blinded observer. Images were flattened using the maximum Z-projection function. Background subtraction was performed, and neuronal processes were thresholded following adjustment of brightness and contrast. Automated Sholl analysis was performed using the ImageJ sholl analysis plugin.

### Excitatory cortical synapse staining

A subset of neurons (2 of 2.7 million) in WT and YAC128 cortical cultures were transfected with a GFP-tagged internally-expressed anti-PSD95 antibody (intrabody) (Gross et al., 2013) at the time of platting. At DIV17 – 19, cells were fixed and stained for VGlut1 and the GluA2 AMPA receptor subunit as previously described (Buren et al., 2016). Briefly, cultures were first live-stained with a primary mouse anti-GluA2 antibody (Millipore), then fixed and stained with a secondary Alexa Fluor 568-conjugated donkey anti-mouse antibody (Invitrogen). Subsequently, cultures were incubated with a primary guinea pig anti-VGlut1 antibody (Millipore), then stained with a secondary AMCA-conjugated donkey anti-guinea pig antibody (Jackson Immuno Research Laboratories). To amplify the GFP fluorescence of the anti-PSD95 intrabody, cultures were also incubated with a primary chicken anti-GFP antibody (1:1000) (Millipore), followed by an secondary Alexa Fluor 488-conjugated antibody (1:500) (Invitrogen). Cultures were imaged on a Zeiss Axiovert 200 M fluorescence microscope (63 x magnification, 1.4 NA), using a Zeiss 702 monochrome camera and Zen software. CPNs expressing the anti-PSD95 intrabody were identified based on their diffuse cytoplasmic GFP fill, with bright GFP-labelled puncta expressed at dendritic spines. A portion of each CPNs arbor, containing multiple secondary and tertiary dendrites, was selected for imaging and sufficient image Z-stacks were acquired to adequately capture all dendritic processes present in a given 63x field. Images were exported to Fiji-ImageJ for analysis by a blinded observer and flattened using the maximum Z-projection function. For each CPN image, the GFP channel was used to identify 3 secondary or tertiary dendritic segments, at least 40 µm away from the CPN soma, over which ROIs were drawn. Following background subtraction, fluorescent puncta in the green (GFP), red (Alexa Fluor 568) and blue (AMCA) channels, visible within dendritic ROIs, were manually thresholded and detected with the analyze particles function. The ImageJ colocalization plugin was used to identify triple-colocalized puncta (PSD95, GluA2 and VGlut1), which we interpreted as functional CPN glutamatergic synapses. Synapse density was defined as the number of triple-colocalized puncta present within a dendritic segment divided by the area of the segment and averaged across all 3 dendrites analyzed in a given CPN.

### Synaptophysin-GCaMP imaging

To directly image cytosolic Ca^2+^ in axonal boutons of neurons in our cortical mono cultures we transfected 1 million cells (of a total 2.7 million) at time of plating with a rat synaptophysin-tagged GCaMP6-M construct (rSyph-GCaMP6m). The rSyph-GCaMP6m construct was created by fusing GCaMP6-M (T.-W. Chen et al., 2013) with the full length rat synaptophysin protein (1 – 307 amino acids) via a small glycine-serine linker and inserting the fused rSyph-GCaMP6-M construct into a pLL3.7-hSyn vector to achieve neuron-selective expression. For some experiments, the same 1 million cells were also transfected at time of platting with an M-cherry-tagged PSD95 construct; in these cases, rSyt-GCaMP6m-expressing presynaptic boutons that colocalized with M-Cherry labeled postsynaptic spines were presumed to be functional synapses.

For all Ca^2+^-imaging experiments, cultures were plated on 8-well cover-glass chambers (Thermo Scientific ™, Nunc ™, Lab-Tek ™) and imaged at DIV (17 - 19) with a Zeiss Axiovert 200 M fluorescence microscope (63 x magnification, 1.4 NA), using a Zeiss 702 monochrome camera and Zen software. Movies were acquired at 10 Hz (100 ms exposure per frame) using the Zen time-series mode with camera-steaming enabled. These experiments were performed in standard ECF (as above) with or without TTX (500 nM) present, but in the absence of PTX.

Spontaneous Ca^2+^ waves were commonly observed in rSyph-GCaMP6m-labelled axons, evident particularly when action potential-dependent Ca^2+^ events were blocked with TTX. Automatically detecting such Ca^2+^ waves proved difficult with algorithms commonly used to quantify activity-related neuronal GCaMP signals. We therefore used the Astrocyte Quantitative Analysis (AQuA) software (running in MATLAB) (Y. Wang et al., 2019) to quantify these waves in the presence of TTX. AQua does not rely on spatially segmenting a Ca^2+^ movie into ROIs based on neuronal morphology, but rather defines events as spatially and temporally connected Ca^2+^ signals surpassing user-defined thresholds. This means spontaneous axonal waves spreading across multiple boutons are typically classified as a single event, because such Ca^2+^-activity is grouped in time and space. Conversely, a given bouton can be involved in multiple axonal waves and thus detected as part of multiple events, as long as the pertinent signals are sufficiently temporally separated. To compare the frequency of aQua-detected spontaneous axonal events between genotypes and under pharmacological manipulations, we counted numbers of events occurring within a standardized area [178.6 μm x 113.1 μm (a maximal 63 x field of view)], during a standardized (3 min) consecutive imaging interval. The same aQua detection parameters, empirically determined to best match a small number of manually analyzed experiments, were applied across all cultures and conditions analyzed to facilitate meaningful comparisons.

In a subset of experiments conducted in the presence of TTX, responses of individual WT or YAC128 rSyph-GCaMP6m-labelled synaptic boutons to caffeine or ionomycin were measured. In these experiments culture fields were imaged for 3 min (as above), after which caffeine (1 mM) or ionomycin (10 μM) was applied and imaging continued for an additional 1-2 min. aQua was not used to analyze these experiments, as an ROI-based algorithm was desired. For caffeine applications, an analyzer (blinded to culture genotype) used Fiji-ImageJ (NIH) to manually assign elliptical ROIs to 10 boutons per movie and exported each ROI’s fluorescence-intensity time-course. Time-courses were subsequently imported to MATLAB, where the curve fitting tool was used to model each time-course’s photobleaching profile based on the initial 3 min recording; this curve was then extrapolated to the entire time-course including caffeine treatment. The resultant “bleaching curve” was subtracted from the raw rSyph-GCaMP6m fluorescence curve and this time-course subsequently divided by the “bleaching curve” to yield a final curve reflecting the caffeine response in DF/F units. Ionomycin experiments were analyzed similarly, except in this case, the blinded analyzer selected 10 active-boutons (showing at least 1 clear Ca^2+^ event during the initial 3 min recording) and 10 inactive-boutons (with no Ca^2+^ events present during the initial 3 min recording). Time-courses derived from inactive-boutons were exported to MATLAB, where the ionomycin-mediated DF/F responses were calculated as above. In the case of active-boutons, spontaneous events occurring during the first 3 min were detected with MATLAB’s findpeaks function and excised from the raw fluorescence time-courses before curve fitting, but otherwise processed as above.

As expected, spontaneous events were far more frequent when rSyph-GCaMP6m -expressing cultures where imaged in the absence of TTX. Most boutons showed many events during a 3 min imagining session. These events, which were presumably driven by the action potential-dependent opening of voltage-gated Ca^2+^ channels, were far shorter in duration than typical TTX-resistant events. To assess for genotype differences in these signals, a blinded analyzer used Fiji-ImageJ to randomly assign elliptical ROIs to 20 boutons per movie and exported each ROI’s fluorescence-intensity time-course to MATLAB. The high frequency of action potential-dependent events often precluded extracting the photobleaching profile of a bouton’s time-course, necessitating an alternative means of calculating the DF/F values of spontaneous events. We therefore averaged the grey-value fluorescence intensity across an entire-time course, subtracted this average fluorescence pointwise from the time-course, then divided pointwise by the average fluorescence to convert to DF/F units. Any nonstationary present in these DF/F time-series due to photobleaching was removed with the MATLAB detrend function. The MATLAB findpeaks function was subsequently used to detect events in these DF/F traces and extract relevant parameters including peak event DF/F amplitudes and event half-amplitude widths. Bouton event frequencies were calculated by dividing numbers of detected events in a DF/F trace by the imaging interval. Bouton event frequencies and event parameters in the absence of TTX were compared between WT and YAC128 cultures under baseline conditions and following application of ryanodine (5 μM).

### iGluSnFR imaging in acute brain slices

Expression of the genetically-encoded intensity-based glutamate-sensing fluorescence reporter (iGluSnFR) (Marvin et al. 2013) in WT and YAC128 mice was achieved with stereotaxic injection of a viral construct as described previously (Parsons et al., 2016). Briefly, under isoflurane anesthesia, 1 – 1.4 μl of the AAV1.hSyn.iGluSnFr.WPRE.SV40 construct (Penn Vector Core; Dr. Loren Looger, Janelia Farm Research Campus of the Howard Hughes Medical Institute) was directly injected into the dorsal striatum of 4 – 6 week old mice. Following surgery, mice were closely monitored for a week to ensure adequate recovery.

After waiting 3 – 6 weeks, to ensure optimal iGluSnFR expression, acute brain slices from 2-3 month-old YAC128 mice and age-matched WT controls were prepared as described previously (Parsons et al., 2016; Koch et al., 2018). Briefly, mice were decapitated following deep isoflurane anesthesia and their brains rapidly removed and placed in an ice-cold slicing solution, bubbled with carbogen (95% O_2_, 5% CO_2_) gas, containing (in mM): 125 NaCl, 2.5 KCl, 25 NaHCO_3_, 1.25 NaH_2_PO_4_, 2.5 MgCl_2_, 0.5 CaCl_2_, and 10 glucose. 300 μm thick striatum-containing sagittal brain slices were cut with a Leica VT1200S vibratome. Slices were subsequently incubated for 30 min in warmed artificial cerebral spinal fluid ACSF containing 2 mM CaCl_2_ and 1 mM MgCl_2_; ACSF constituents and concentrations were otherwise identical to the slicing solution (above).

Slices were transferred to a submerged recording chamber for experiments and perfused with carbogen-bubbled ACSF at a rate of 2 – 3 ml/min at room temperature. Cortical release of glutamate into the striatum was evoked by delivering paired 0.1 ms electrical pulses at 100 Hz with an A-M Systems isolated pulse stimulator (Model 2100) and tungsten monopolar stimulating electrode (tip resistance - 0.1 MΩ). The electrode was placed into a corpus callosum segment adjacent to the dorsal striatum at an approximate 50 – 100 μm depth. During and immediately prior to the electrical stimulation, iGluSnFR fluorescence was excited with a 470 nm LED; slices were not illuminated between experimental measurements to minimize phototoxicity and bleaching. Stimulation and LED activation were triggered by Clampex software (Molecular Devices, Sunnyvale, CA). iGluSnFR fluorescence was isolated with a 530 nm bandpass filter and imaged with a CCD camera (1 M60, Pantera, Dalsa) and XCAP software (Epix, inc.) at 150 Hz with 8 × 8-pixel binning. Experimental measurements encompassing four stimulation trials and two blank trial were performed at 3-minute intervals. The four stimulation trials were averaged and the blank trials, in which slice-fluorescence was imaged without electrical stimulation, were averaged and used to account for photobleaching and to calculate the stimulation mediated changes in iGluSnFR fluorescence over basal fluorescence (ΔF/F) as described previously (Parsons et al., 2016). Videos were analyzed offline with ImageJ software. The average iGluSnFR signal was measured over 10 × 10-pixel (93.8 × 93.8 μM) region of interested (ROI) placed over the maximal area of evoked iGSnFR activity within the striatum adjacent to the stimulating electrode.

### Experimental design and statistical analysis

Statistical analysis and creation of figures was performed using GraphPad Prism (version 7). All data distributions were tested for normality with the D’Agostino-Pearson omibus normality test.

The Student’s unpaired t-test was used for unpaired comparisons between two data groups, such as when mean mEPSC frequencies were compared between WT and YAC128 neurons, as long both data groups were normally distributed. When one or both groups failed the D’Agostino-Pearson omibus normality test, the non-parametric Mann Whitney test was used instead.

When parameters of the same group of neurons or axonal boutons were compared before and after a drug treatment, statistical significance was assessed with the Student’s paired t-test, unless data points in the control or drug-treatment group failed the D’Agostino-Pearson omibus normality test; in which case, the non-parametric Wilcoxon matched-pairs signed rank test was used instead.

A two-way ANOVA with the Bonferroni post-test was used when testing for genotype differences in a dependent variable measured at different time points, as was the case for our brain-slice iGluSnFR experiments.

Depending on the experimental design, “n” numbers in figures refer either to number of neurons, numbers of imaged culture fields, numbers of individual axonal boutons, or numbers of brain slices. Clarifying details are present within individual figure legends, as are the numbers of culture batches or mice used.

Differences in mean values were considered significant at p<0.05, and significance levels are indicated in figures as follows: *p<0.05, **p<0.01, ***p<0.001, ***p<0.0001. Comprehensive descriptions of statistical analysis are included in figure legends.

## Acknowledgments

We thank Dr. Anne Marie Craig (UBC) for expert advice on experiments using rSyph-GCaMP6m, and Dr. Lily Zhang and Dr. Rujun Kang for technical support and assistance. The work was supported by funding from the Canadian Institutes of Health Research (CIHR) Foundation grants FDN-143210 to LAR and FDN-154278 to MRH. CB was supported by a University of British Columbia 4-year Graduate Fellowship (UBC 4-YF). ASD was supported by a CIHR Canada Graduate Scholarship Doctoral award and a UBC 4-YF. EK was supported by a CIHR Canada Graduate Scholarship Master’s award and UBC 4-YF. MES was supported by a Vanier Canada Graduate Scholarship and a UBC 4-YF. WN was supported by a UBC-CIHR-MD/PhD studentship and a Vanier Canada Graduate Scholarship. LAR holds the UBC Department of Psychiatry Louise A. Brown Chair in Neuroscience. MRH holds a Canada Research Chair.

## Figure legends

**Supplemental Fig 1:**
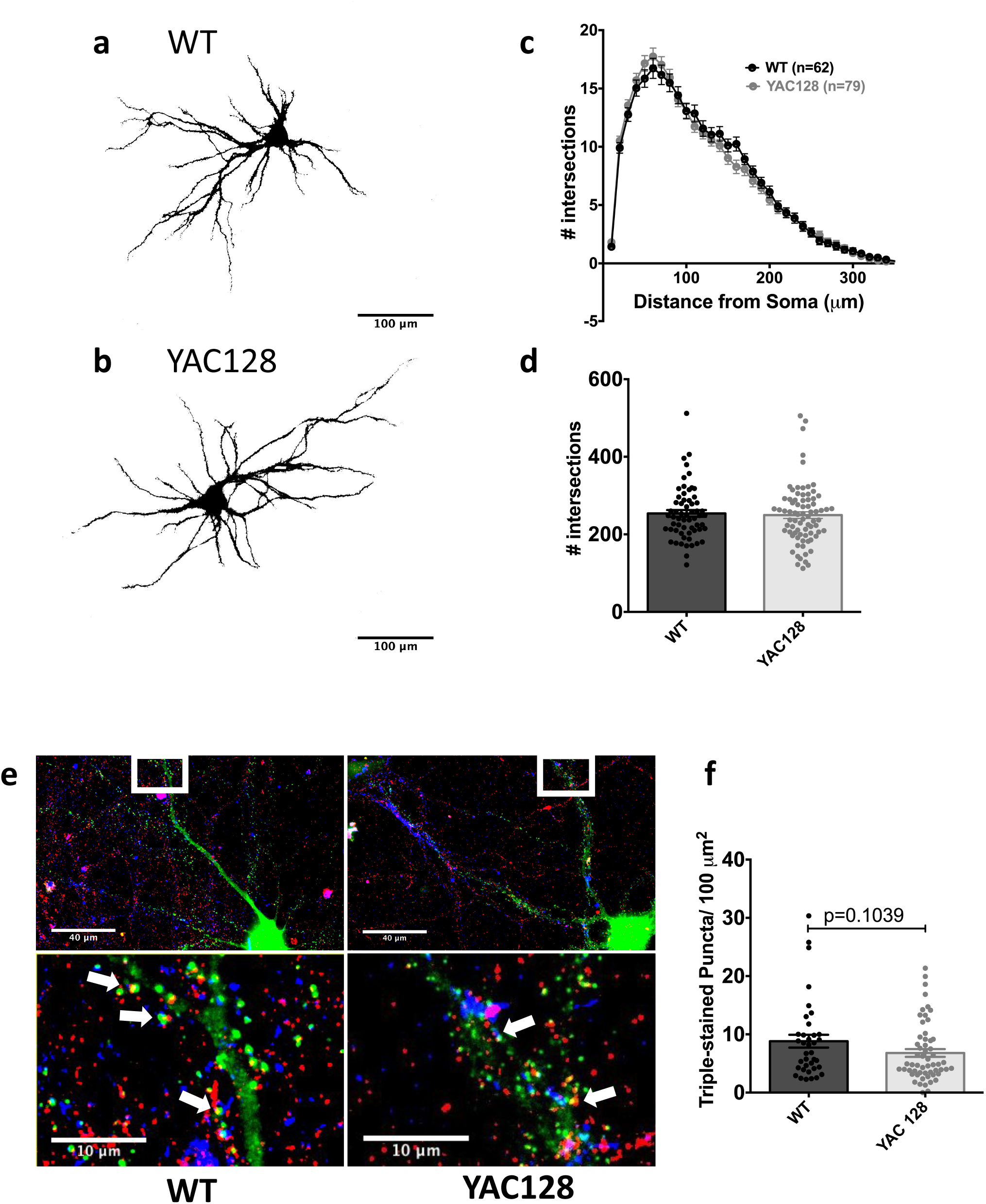
Dendritic complexity and excitatory synapse numbers are similar in WT and YAC128 cortical cultures. **A, B**. Representative images generated by thresholding a merged Z-stack of a green fluorescent protein (GFP)-expressing WT CPN (**A**) and YAC128 CPN (**B**). **C**. Number of dendritic intersections through concentric sholl circles centered on the soma of GFP-filled WT and YAC128 CPNs of radii between 10 μm and 450 μm. WT and YAC128 CPNs have nearly identical dendritic branching distributions [2-way ANOVA; Genotype: F(1, 6165)= 1.168, p<0.2799; Distance from soma: F(44, 6165)=381.6, p<0.0001; Interaction: F(44, 6165)=0.9868, p<0.4966]. **D**. Total numbers of sholl intersections, which reflect general neuronal complexity, were similar in WT and YAC128 CPNs: 254.0 ± 8.8 (n=61; 4 cultures) and 249.5 ± 8.7 (n=78; 4 cultures) respectively [p=0.7263 (exact); Mann Whitney test]. **E**. A representative merged Z-stack from a WT CPN (left panels) and YAC128 CPN (right panels) with merged green, red and blue channels showing staining of a genetically-encoded internally- expressed GFP-tagged anti-PSD-95 antibody (green), anti AMPA receptor GluA2 subunit immuno-staining (red) and anti VGlut1 immuno-staining (blue) respectively. The bottom panel shows an expanded view of a segment of dendrite with arrows pointing to a subset of co-localized PSD-95, GluA2 and VGlut1 puncta, which were counted as presumed functional synapses. Note, for illustrative purposes the brightness and contrast of the individual channels were adjusted to best illustrate the punctate fluorescence. **F**. Numbers of presumed functional synapses (identified as above) per 100 μm^2^ of secondary and tertiary dendrites of WT and YAC128 CPNs: 8.8 ± 1.1 (n=37; 2 cultures) and 6.8 ± 0.7 (n=57; 3 cultures), respectively. Although synapse numbers were lower in YAC128 cultures this did not reach statistical significance [p=0.1039 (exact); Mann Whitney test].

**Supplemental Fig 2:**
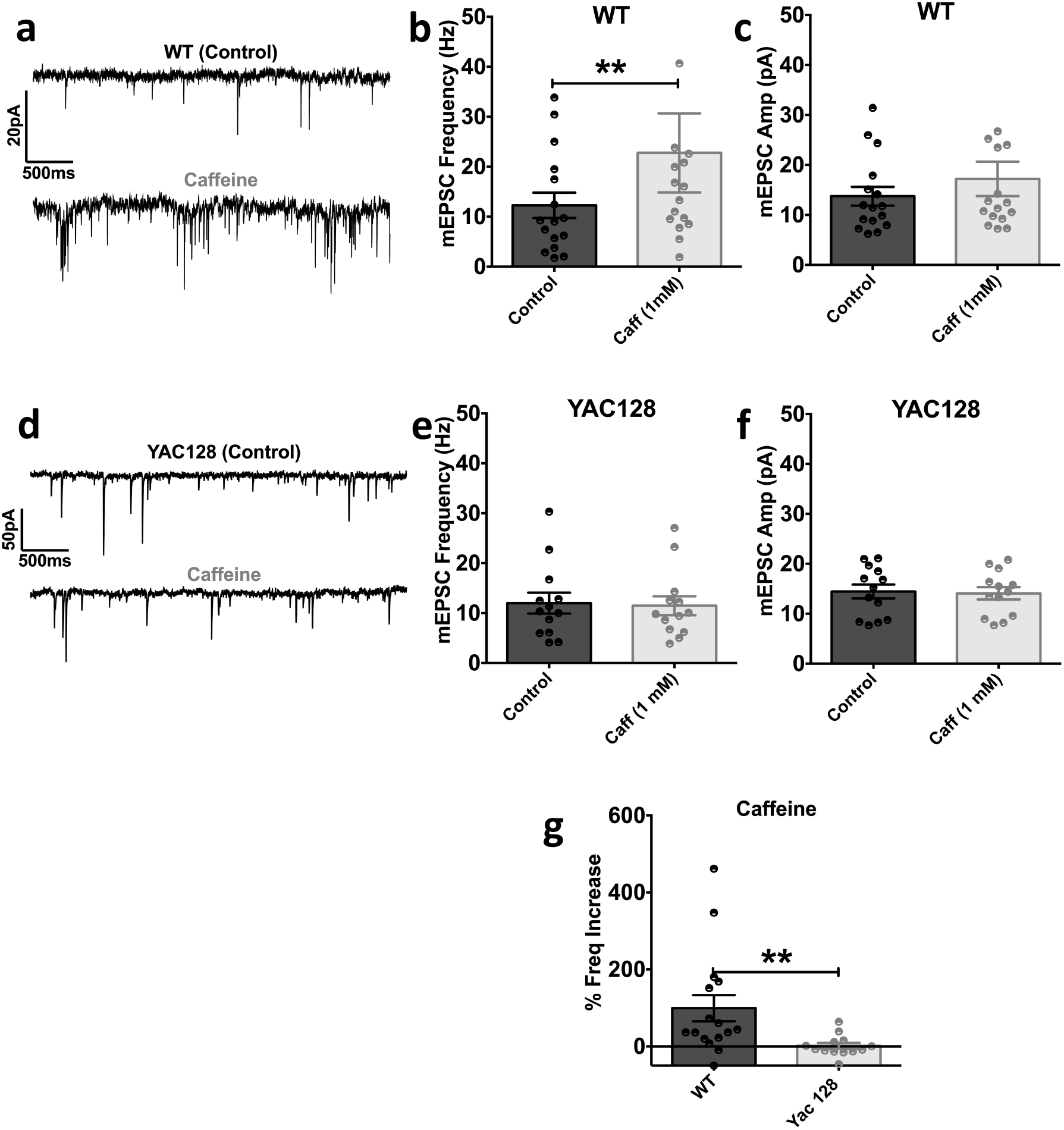
Releasing ER Ca^2+^ with caffeine increases the CPN mEPSC frequency in WT, but not YAC128, CPNs. Voltage clamp recordings were made from cultured WT and YAC128 CPNs at -70 mV in the presence of TTX (500 nM) and PTX (50 μM), under control conditions and during subsequent local caffeine (1 mM) application. **A, D:** Representative traces from WT (A) and YAC128 (D) CPNs, under control (top traces) and caffeine (bottom traces) conditions. In this CPN, caffeine (1 mM) substantially increased the mEPSC frequency. **B**. Caffeine (1 mM) significantly increased the mean mEPSC frequency in WT cultured CPNs from 12.3 ± 2.5 Hz to 22.8 ± 8.0 Hz, [p=0.0076 (exact); n=16; 8 cultures; Wilcoxon matched-pairs signed rank test]. **C**. Caffeine (1 mM) did not significantly affect the mean WT CPN mEPSC amplitude (control: 13.7 ± 1.9 pA, caffeine: 17.2 ± 3.4 pA), [p=0.8603 (exact); n=16; 8 cultures; Wilcoxon matched-pairs signed rank test]. **E**. In YAC128 CPNs, caffeine (1 mM) did not significantly affect the mean CPN mEPSC frequency (control: 12.0 ± 2.1 Hz, caffeine: 11.5 ± 1.9 Hz), [t(12)=0.4808; p=0.6393; n=13; 5 cultures; Student’s paired-t test]. **F**. In YAC128 CPNs, caffeine (1 mM) also did not significantly affect the mean CPN mEPSC amplitude (control: 14.5 ± 1.4 pA, caffeine: 14.1 ± 1.2 pA), [t(12)=0.8086; p=0.4345; n=13; 5 cultures; Student’s paired-t test]. **G**. Percent change in mEPSC frequency mediated by Caffeine (1 mM) application to WT and YAC128 CPNs (calculated from the responses in panels **B** and **E**). Caffeine (1 mM) elicited a significantly greater increase in the mEPSC frequency in WT, compared to YAC128, CPNs [WT: 99.2 ± 34.1 % (n=16; 8 cultures), YAC128: 1.2 ± 7.6 % (n=13; 5 cultures)], [p=0.0025 (exact); Mann Whitney test].

